# Two heads are better than one: Single stranded DNA translocation of UvrD-family dimers vs. monomers

**DOI:** 10.64898/2026.08.07.743547

**Authors:** Kacey N. Mersch, Binh Nguyen, Alexander G. Kozlov, Timothy M. Lohman

## Abstract

UvrD-family Superfamily 1A helicases are processive ATP-dependent motor proteins that function during DNA replication, recombination, repair, and transcription. UvrD-family monomers translocate along single stranded (ss) DNA with 3’-to-5’ directionality but must be activated by dimerization to become helicases in the absence of force or accessory factors. *Mycobacterium tuberculosis* (*Mtb*) UvrD1 helicase forms dimers via a disulfide bond between native cysteines in the 2B sub-domains of each monomer. *E. coli* UvrD forms non-covalent dimers using the same 2B domain interface as in *Mtb* UvrD1. Using both ensemble and single DNA molecule approaches we examined an *E. coli* UvrD variant (R421C), which forms covalent dimers with constitutive helicase activity. For the first time this has enabled us to compare the ssDNA translocation and helicase activities of covalent dimers and monomers. Crosslinked UvrD dimers exhibit much higher ssDNA translocation processivities than monomers, although with similar translocation rates. Crosslinked UvrD dimers also show highly processive DNA unwinding of thousands of base pairs, much higher than non-crosslinked UvrD dimers, while monomers show no DNA unwinding activity. DNA unwinding rates of crosslinked UvrD dimers are only ∼20% slower than ssDNA translocation rates, indicating they are “active” helicases that directly facilitate duplex destabilization.

## Introduction

Helicases are nucleic acid motor proteins that use ATP binding/hydrolysis to catalyze the unwinding (strand separation) of duplex DNA and function in nearly all aspects of genome maintenance[2–6]. Processive helicases also translocate directionally along single stranded (ss) DNA[7–9]. Helicases/translocases are divided into six superfamilies (SF1-6), with SF3-6 enzymes forming hexamers, whereas SF1 and SF2 are non-hexameric[10, 11]. SF1 family members are further defined by the direction of translocation along ssDNA. SF1A members translocate with 3’-to-5’ polarity on ssDNA, while SF1B members translocate with 5’-to-3’ polarity[6]. *E. coli* (*Ec*) UvrD [12, 13] is an SF1A helicase/translocase that functions in methyl-directed mismatch DNA repair[14], nucleotide excision repair[15], replication restart[16, 17], recombination[18, 19] and transcriptional control[20]. *Ec* UvrD is structurally similar to *Ec* Rep[21], *Bacillus stearothermophilus (Bst)* PcrA[22, 23], and the homologous *Mycobacterium tuberculosis* (*Mtb*) UvrD1[11]. SF1A monomers are composed of four sub- domains (1A, 1B, 2A, 2B). The 1A and 2A sub-domains are the motor domains that bind ATP at their interface. The 2B sub-domain is rotationally dynamic and plays an important regulatory role[24]. While monomers of UvrD, Rep, PcrA, and UvrD1 are rapid and processive ssDNA translocases, they have no detectable helicase activity [9, 24–29]. Monomer helicase activity is prevented due to an auto- inhibitory interaction of the 2B sub-domain with duplex DNA[1, 24]. In the absence of assisting force applied to the DNA[30–32], this auto-inhibition can be relieved by deletion of the 2B sub-domain[28, 31, 33], by dimerization [1, 26, 27, 29, 34–49], through interactions with accessory proteins (MutL with UvrD[50, 51], PriC with Rep[52], and RepD with PcrA[53]), or by crosslinking of the 2B sub-domain within the monomer [54].

Recent structures of the *Mtb* UvrD1 dimer bound to a ss/dsDNA junction **(Figure 1A)** show the structural basis for helicase activation by dimerization [1, 24]. The UvrD1 dimer is stabilized by a native disulfide crosslink formed between the same cysteine (Cys451) within the 2B domains of each UvrD1 subunit[1, 29] and thus differs from the non-covalent dimers formed by *Ec* UvrD, *Ec* Rep and *Bst* PcrA[24]. Importantly, dimerization via the 2B sub-domains prevents the 2B sub-domain of the lead subunit from forming the auto-inhibitory interaction with the duplex DNA that occurs within the monomer [1, 24]. However, dimerization of *Ec* UvrD involves the same 2B-2B interface as observed for *Mtb* UvrD1. In fact, a variant of *Ec* UvrD (R421C) in which a cysteine is placed within the 2B sub-domain in a structurally homologous position to UvrD1 C451, can form a redox-dependent crosslinked dimer with constitutive helicase activity [1]. Recent studies showed that *Ec* Rep and *Bst* PcrA can also form crosslinked constitutively active dimeric helicases via the same 2B domain interface [55]. Hence, all four of these SF1A enzymes can be activated via a conserved mechanism involving dimerization via the 2B sub-domains [1, 24, 55].

**Figure 1.**
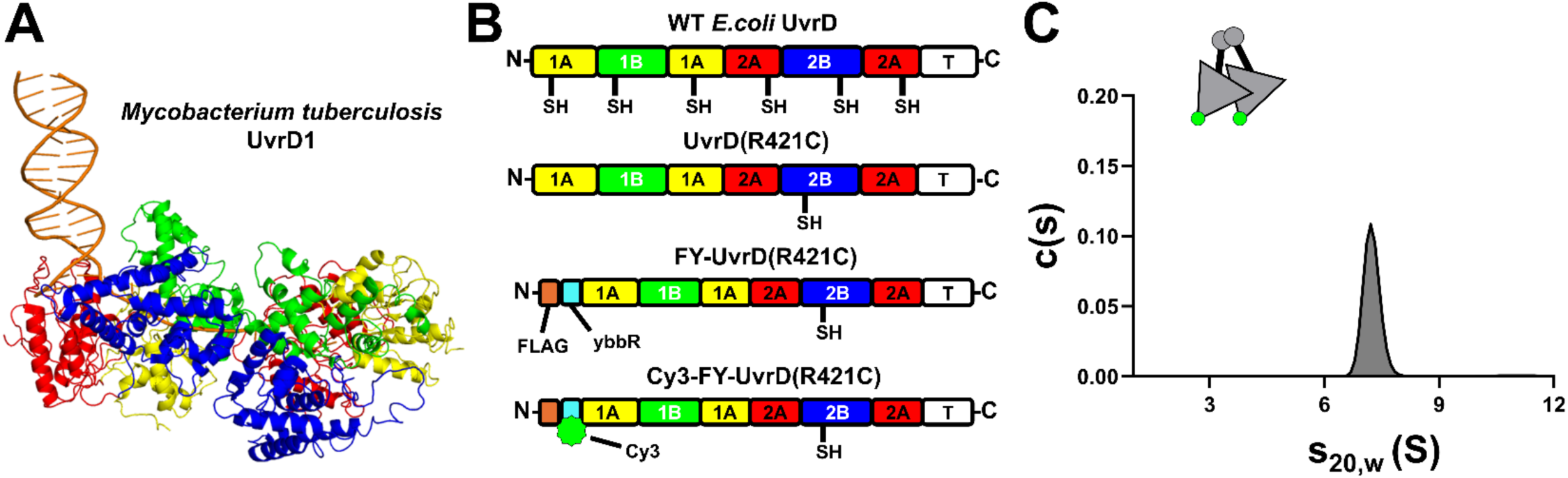
UvrD protein constructs. **(A)** Structure of dimeric *Mtb* UvrD1 bound to a 3’-ss/ds DNA junction (PDB ID: 9DES) [1]: DNA (orange), 1A domain (yellow), 2A domain (red), 1B domain (green), 2B domain (blue), tudor domain (T) (unresolved in the structure). **(B)** Domain structure of wt *E. coli* UvrD, UvrD(R421C), FLAG-ybbR-UvrD(R421C) (FY- UvrD(R421C), and Cy3- FY-UvrD(R421C). **(C)** Sedimentation velocity experiment of purified crosslinked Cy3-FY- UvrD(R421C) shows a single c(s) peak at s_20,w_=7.2 S corresponding to a dimer in Buffer T20-20.

It has been difficult to study non-covalent *Ec* UvrD dimers since dimers readily dissociate into monomers at low concentrations and can also forms tetramers at the higher concentrations needed to form dimers [56]. However, our ability to purify crosslinked, constitutively active *Ec* UvrD(R421C) dimers allows us to directly compare the activities of *Ec* UvrD monomers and crosslinked dimers at the same concentrations and under the same solution conditions. Our single DNA molecule studies show that a single crosslinked UvrD dimer can unwind thousands of base pairs of DNA and can translocate along ssDNA for thousands of nucleotides with ∼two-fold higher processivity than monomers, although they exhibit similar translocation rates. These studies indicate that covalent dimerization of SF1A enzymes forms constitutively active helicases with high processivity. Comparison of the ssDNA translocation and DNA unwinding rates of UvrD dimers indicate that they play a direct role in destabilizing the DNA duplex and thus function as active DNA helicases [2, 38, 57, 58].

## Results

### Purification of Crosslinked FY-UvrD(R421C) Dimers

The *E. coIi (Ec)* UvrD constructs used in this study are shown in **Figure 1B**. In UvrD(R421C) the six native cysteines present in wild type (wt) *Ec* UvrD were replaced with serine [13, 55, 59] and the native arginine at position 421 with cysteine as described [1, 55]. In FY-UvrD(R421C), a FLAG and ybbr-tag (FY) were also placed in tandem on the N-terminus of UvrD(R421C). We expressed and purified these proteins as described [13, 37]. We then labeled FY-UvrD(R421C) with a Cy3 fluorophore using Surfactin phosphopantetheinyl transferase (Sfp) as described [60, 61] (see **Materials and Methods**) to form Cy3-FY-UvrD(R421C).

In the absence of reducing agent, UvrD(R421C) an FY-UvrD(R421C) form crosslinked dimers [1, 55]. During purification of FY-UvrD(R421C), the monomeric species is largely removed and yields 96% cross-linked dimer on average as determined by sedimentation velocity experiments (**Supplemental Figure 1)**. The sedimentation coefficient of UvrD monomers is s_20,w_ = 4.6 ± 0.2 S, and UvrD dimers is s_20,w_ = 7.4 ± 0.2 S [1, 55]. Further purification during the labeling of FY-UvrD(R421C) with Cy3 for single-molecule studies leaves us with a homogeneous population of crosslinked dimers as shown by sedimentation velocity experiments, displaying a single peak at s_20,w_ = 7.2 S, shown in Figure 1C. The position of the c(s) peaks are independent of protein concentration indicating that the dime do not dissociate into monomers in the absence of reducing agent. However, upon addition of 1mM dithiothreitol (DTT), protein dimers are converted fully to monomers (s_20,w_ = 4.6 S) **(Supplemental Figure 1)**. Under the same conditions and protein concentrations wt *Ec* UvrD exists only as monomers [56].

### ssDNA Translocation

We first examined ssDNA translocation of both the monomeric and dimeric crosslinked FY- UvrD(R421C) using an ensemble stopped-flow fluorescence assay as described [8, 9, 28, 47, 62, 63]. The assay monitors translocation of FY-UvrD(R421C) on a series of ss oligodeoxythymidylate molecules, (dT)_L_, of various lengths (L=45, 55, 79, 104 and 124 nucleotides) with a Cy3 fluorophore at the 5’ end as depicted in **Figure 2A and 2B**. Protein is pre-bound to the Cy3-(dT)_n_ in Buffer T20-20 in one syringe of the stopped-flow and then mixed with ATP and heparin in Buffer T20-20 in the other syringe. The pre-bound enzymes translocate in the 3’-to-5’ direction and upon reaching the 5’-end interact with the Cy3 fluorophore resulting in an enhancement of Cy3 fluorescence due to PIFE (Photoisomerization-related Fluorescence Enhancement) [64–66]. The presence of heparin serves as a trap to bind any FY-UvrD(R421C) that dissociates from the Cy3-(dT)_n_ thus ensuring single round conditions [9, 67]. The monomeric FY-UvrD(R421C) was prepared by adding 1mM DTT to the buffer containing dimeric FY-UvrD(R421C).

**Figure 2.**
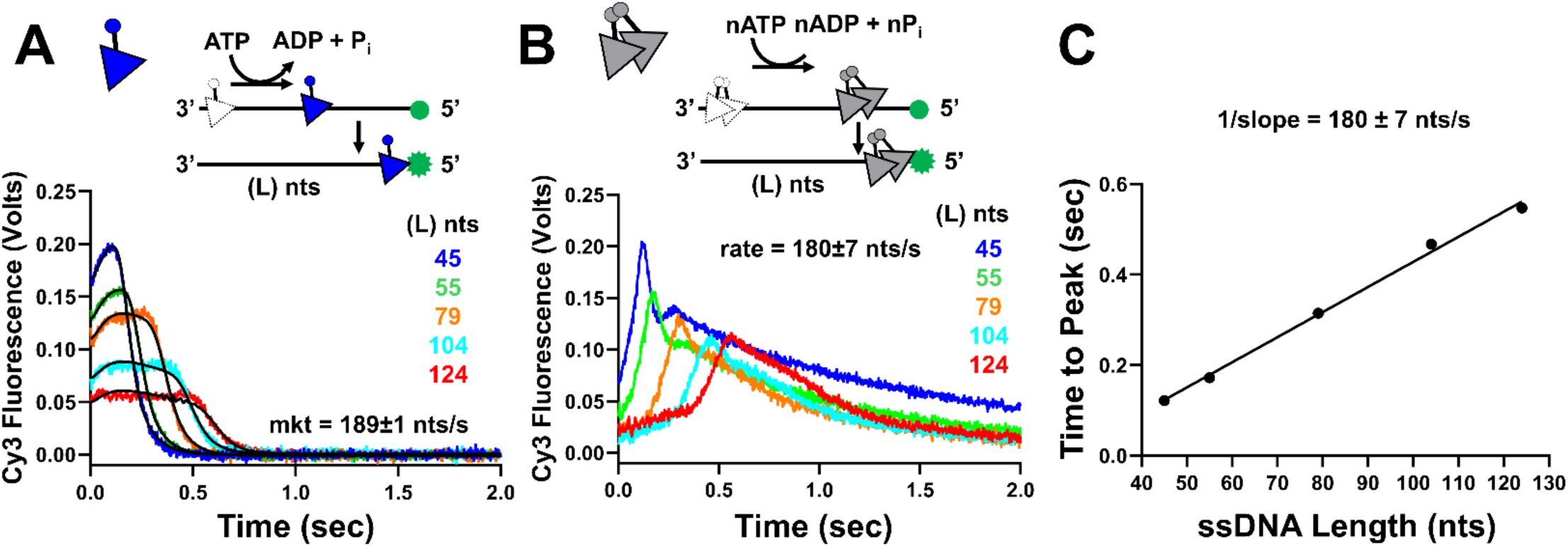
Stopped-flow ssDNA translocation of FY-UvrD(R421C) dimers vs. monomers. **(A)** Single-round stopped-flow fluorescence ssDNA translocation time courses for FY-UvrD(R421C) monomers (**A**) on a series of 5’-Cy3-(dT)_L_ (L= 45, 55, 79, 104, 124 nucleotides) in Buffer T20-20, 500 μM ATP, 1 mM MgCl_2_ and 4mg/mL heparin (post-mix) plus 1mM DTT, 25 °C (50 nM monomer and 100 nM DNA). Continuous black lines show the best global fit of the five time courses to Scheme 1 ((Eq. (S3)) yielding a macroscopic ssDNA translocation rate of 189±1 nt/sec (Table 1). **(B)** Single-round stopped-flow fluorescence ssDNA translocation time courses for FY-UvrD(R421C) dimer translocation on a series of 5’- Cy3-(dT)_L_ (L= 45, 55, 79, 104, 124 nucleotides) in Buffer T20-20, 500μM ATP, 1mM MgCl_2_ and 4mg/mL heparin (post-mix). **(C)** Time-to-peak analysis of the data in panel B yields a ssDNA translocation rate (1/slope = 180 ± 7 nt/sec) (Table 1).

**Table 1:** Rates and processivities of ssDNA translocation by FY-UvrD(R421C) and Cy3-FY-UvrD(R421C)

| Rate (nts/sec) | <N>, average nts translocated | Experiment | Monomer or Dimer | Tension pN | Labeled with Cy3 | [ATP] μM |
| --- | --- | --- | --- | --- | --- | --- |
| 189±1 | | Stopped-Flow | Monomer | | No | 500 |
| 180±7 | | Stopped-Flow | Dimer | | No | 500 |
| 184±94 | 1100±20 | Optical Tweezers | Monomer | 10 | Yes | 500 |
| 200±77 | 2309±22 | Optical Tweezers | Dimer | 10 | Yes | 500 |
| 191±89 | 1208±18 | Optical Tweezers | Monomer | 20 | Yes | 500 |
| 190±88 | 2841±84 | Optical Tweezers | Dimer | 20 | Yes | 500 |

The stopped-flow ssDNA translocation time courses for the monomeric FY-UvrD(R421C) **(Figure 2A)** are similar to those observed previously for wt *Ec* UvrD monomers and indicate random initial binding of monomers to the Cy3-(dT)_L_ [9, 67–70]. The time courses were globally fit to **Scheme 1** (**Eq. (3)**) yielding a macroscopic translocation rate of 189 ± 1 nt/s (Table 1), the same as measured previously for wild type UvrD monomers under the same conditions (193 ± 4 nt/s) [9, 67, 68]. In contrast, the time courses for ssDNA translocation of the crosslinked FY-UvrD(R421C) dimers were qualitatively different and showed a sharp peak in Cy3 fluorescence that moved to longer times as the Cy3-(dT)_L_ length increased **(Figure 2B)**. These qualitatively different time courses suggest that some fraction of the crosslinked FY-UvrD(R421C) dimers initiate at the same position on each Cy3-(dT)_L_ (likely the 3’- end) rather than randomly along the ssDNA as for the monomer (**Figure 2A**) [70]. We analyzed the dimer time courses using a “time-to-peak” analysis performed by plotting the time at which the acute peak occurs as a function of ssDNA length and fit the data to a straight line to obtain the macroscopic translocation rate (**Figure 2C**), as described [62, 70]. This analysis assumes that the dimers all initiate translocation from the 3’-end and yields a ssDNA translocation rate of 180 ± 7 nt/s (Table 1), which is the same within error as the monomer translocation rate.

**Scheme 1.**
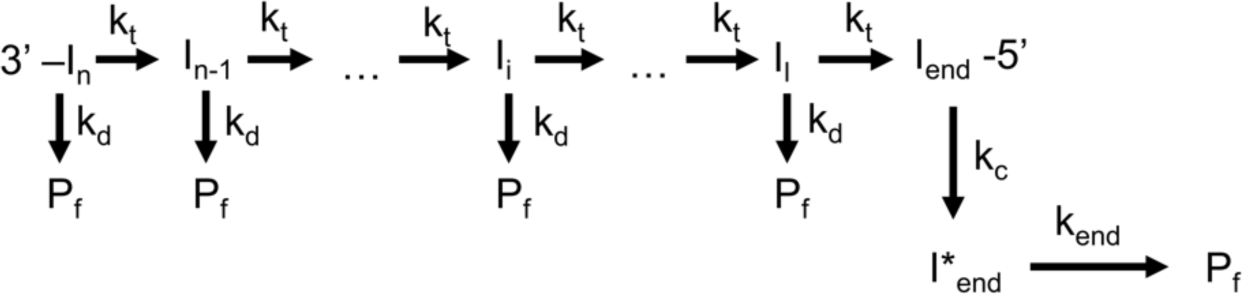

### Single molecule experiments reveal crosslinked Cy3-FY-UvrD(R421C) dimers translocate at the same rate as monomers, but with much higher processivity

The results from the stopped-flow experiments suggest that UvrD dimers and monomers translocate along ssDNA with the same macroscopic rate. To examine this further, we used a single DNA molecule optical tweezers experiment with fluorescently labeled protein to monitor ssDNA translocation directly. In addition to rates, these experiments also allow us to obtain a quantitative estimate of translocation processivity. Our ability to purify fully crosslinked UvrD dimers was essential for unambiguous interpretation of the single molecule experiments. Any sample of non-covalent UvrD dimers will also contain monomers and dimer dissociation to form monomers will occur during translocation making it impossible to identify monomers vs. dimers.

The optical tweezer experiments were performed using a LUMICKS C-trap depicted in **Figure 3A** (see also **Supplemental Figure 2A)**. A single dsDNA molecule (20,425 base pairs) possessing a 5x biotin tag on both the 3’- and 5’- ends of one DNA strand was secured between two streptavidin-coated polystyrene beads held within each of the two optical traps in PBS buffer. Tension was placed on the dsDNA to induce melting of the DNA and dissociation of the non-tethered DNA strand, leaving a fully ssDNA (20,425nts) tethered between the two beads (**Figure 3A).** The ssDNA was then moved into a channel of the flow cell containing Cy3 labeled protein in Single Molecule Imaging Buffer containing 0.5 mM ATP and a constant tension (10 or 20pN) was applied to the ssDNA using a force-clamp. Kymographs were obtained by repeatedly scanning the ssDNA while exciting the Cy3-labeled protein with a 532nm laser. Representative kymographs show ssDNA translocation of Cy3-FY-UvrD(R421C) monomers in the presence of 1mM DTT **(Figure 3B)** and crosslinked FY-UvrD(R421C) dimers **(Figure 3C)**. Single molecule trajectories were then created by tracking the positions of the translocating proteins along the ssDNA. None of the trajectories showed significant pausing or spontaneous changes in rate, hence we fit the trajectories with linear lines to determine ssDNA translocation rates.

**Figure 3.**
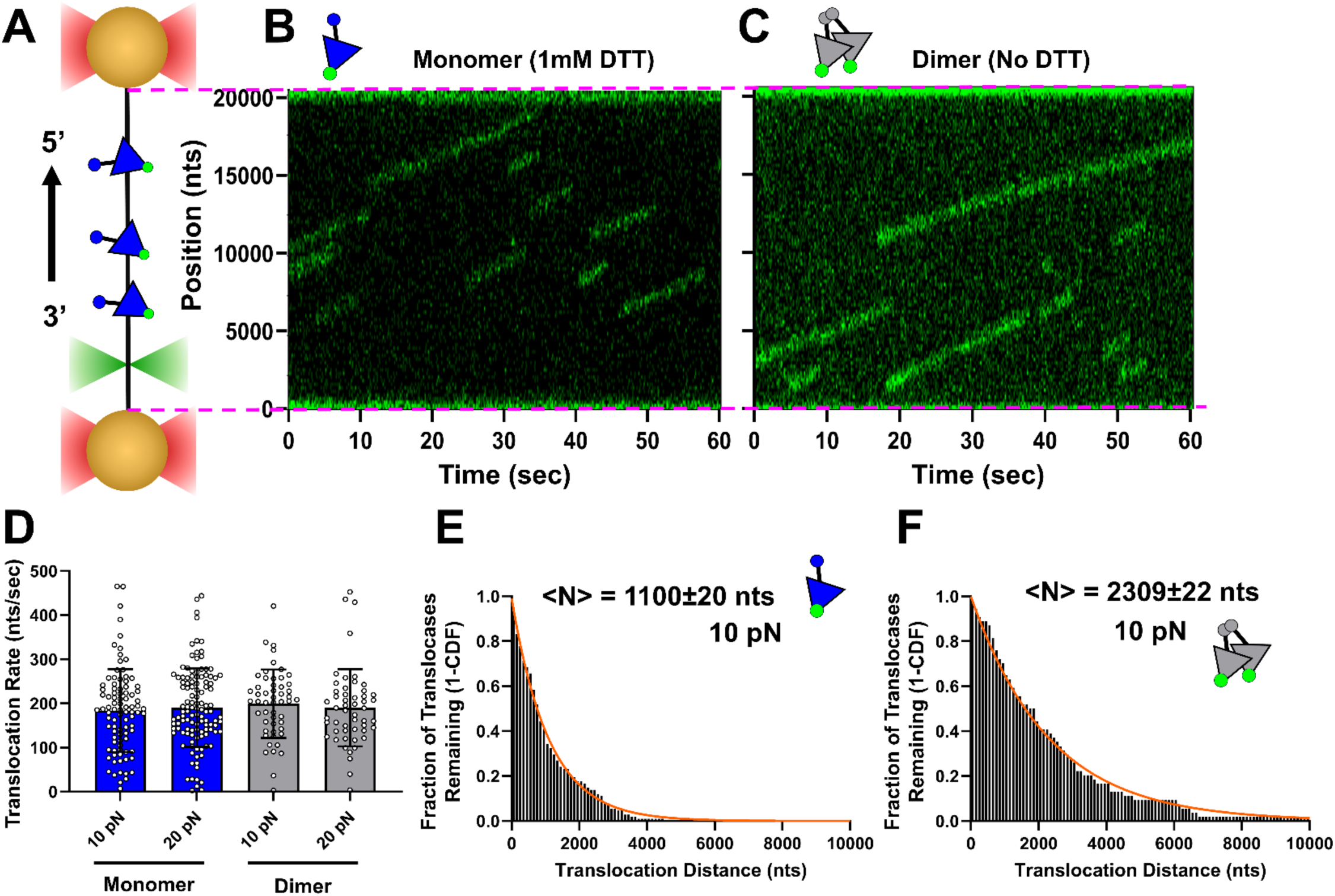
Single DNA molecule experiments show Cy3-FY-UvrD(R421C) monomers and dimers translocate at the same rate, but dimers possess higher processivity. **(A)** Schematic of the optical tweezer (LUMICKS) experiment. **(B)** Kymographs showing ssDNA translocation of Cy3-FY-UvrD(R421C) monomers in Single Molecule Imaging Buffer plus 1 mM DTT, 25 °C. **(C)** Kymographs showing ssDNA translocation of crosslinked Cy3-FY-UvrD(R421C) dimers in Single Molecule Imaging Buffer (No DTT), 25 °C. **(D)** Individual translocation rates at 10pN: monomer - 184 ± 94 nt/s, dimer - 200 ± 77 nt/s; at 20 pN, monomer - 191 ± 89 nt/s, dimer - 190 ± 88 nt/s (mean ± S.D.) (Table 1). Fraction of monomers **(E)** and dimers **(F)** remaining bound vs. distance travelled (in nucleotides) and the fit to Eq. (5) to obtain the average processivity, <N>, at 10 pN (Table 1).

**Figure 3D** and Table 1 summarize the translocation rates for Cy3-FY-UvrD(R421C) monomers and crosslinked dimers on ssDNA held at forces of 10 pN and 20 pN. At 10pN monomers translocate with rates of 184 ± 94 nucleotides (nt)/s (Table 1), and dimers with rates of 200 ± 77 nt/s (Table 1). At 20pN monomers translocate with rates of 191 ± 89 nt/s (Table 1), and dimers with rates of 190 ± 88 nt/s (Table 1). Therefore, both monomers and crosslinked dimers exhibit the same macroscopic ssDNA translocation rates at 0.5 mM ATP, independent of ssDNA tension, consistent with the stopped-flow experiments.

To assess ssDNA translocation processivity we measured the distance that each enzyme translocated (run length) and plotted the fraction of translocases remaining bound vs. translocation distance for monomers (**Figure 3E**) and crosslinked dimers (**Figure 3F**). These plots were well described by single exponential decays and were fit to **Eq. (5)** to obtain the average number of nucleotides translocated [2, 71, 72] summarized in Table 1. At 10 pN, <N> = 1100 ± 20 nt (Table 1) for the monomers, and <N> = 2309 ± 22 nt (Table 1) for the dimers. At 20pN the average number of nucleotides translocated increased slightly for both monomers (<N> = 1208 ± 18 nt) (Table 1) and dimers (<N> = 2841 ± 84 nt) (Table 1) **(Figure S3**), but still showed a more than two-fold higher processivity for the dimers. Therefore, although monomers and dimers translocate along ssDNA at the same average rates, crosslinked dimers translocate ∼two-fold further than monomers on average.

### Crosslinked UvrD dimers have higher DNA unwinding processivity than non-covalent UvrD dimers

We next used an all-or-none fluorescence stopped-flow assay [55] to examine the DNA unwinding (helicase) activities of crosslinked FY-UvrD(R421C) dimers and FY-UvrD(R421C) monomers. The DNA substrates used are depicted in **Figure 4A** and contain a 3’-(dT)_20_ loading site and a variable duplex DNA length of 18, 21, 25, 40, or 50 base pairs. The 5’-ended strand at the blunt end of the duplex DNA was labeled with a Cy5 fluorophore and the 3’-ended strand was labeled with a black hole quencher 2 (BHQ2). DNA unwinding was monitored by the increase in Cy5 fluorescence that occurs when the duplex DNA strands are completely separated [51]. Protein and DNA substrate are pre-bound in one syringe of the stopped-flow in Buffer T20-20 and then mixed with ATP (0.5 mM final concentration) and an excess of unlabeled DNA in Buffer T20-20 in the other syringe. The unlabeled DNA serves as a trap to bind free protein to prevent re-initiation of unwinding of the labeled DNA substrate, ensuring only a single round of DNA unwinding [29, 50, 51].

**Figure 4.**
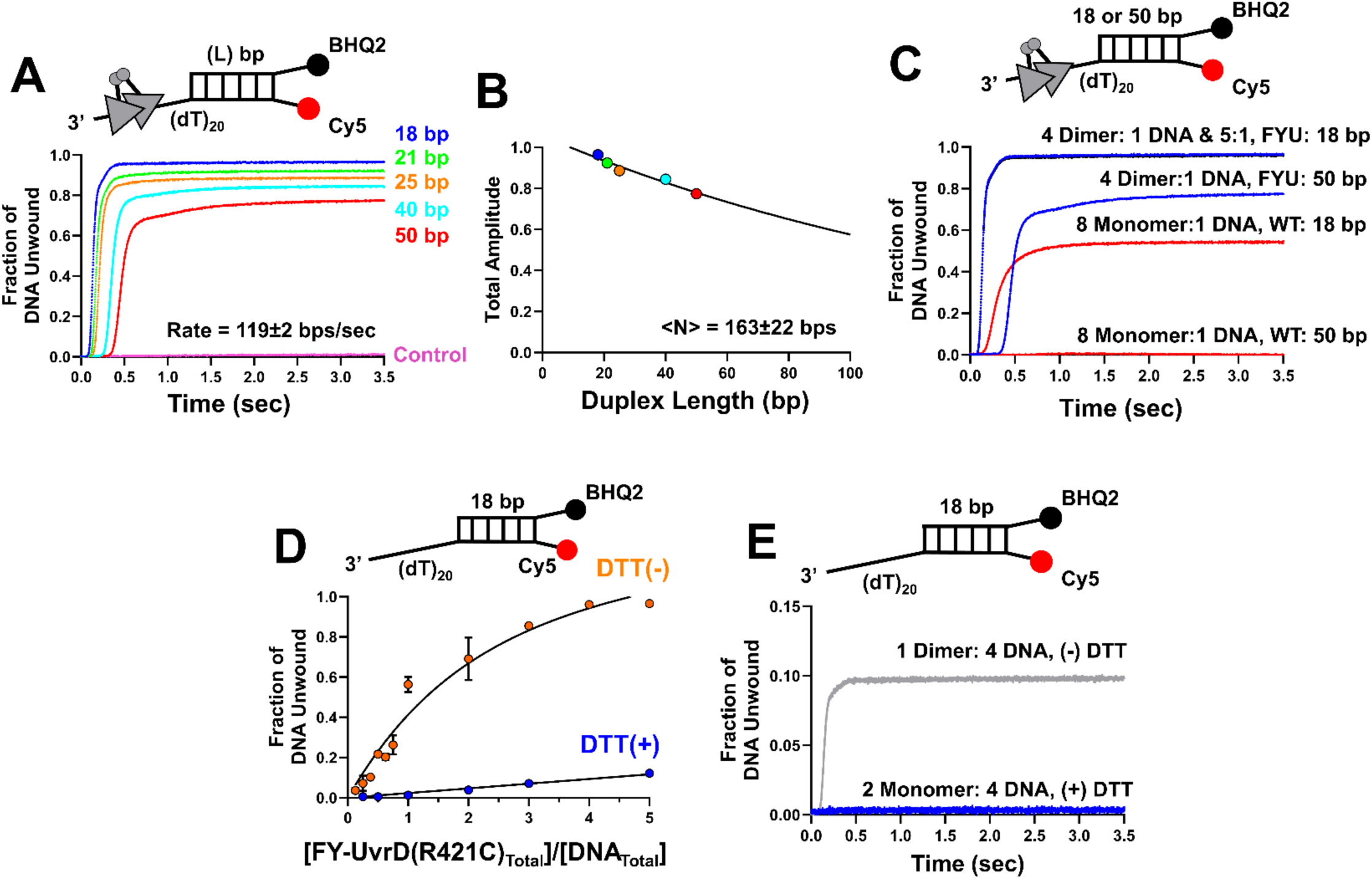
Crosslinked FY-UvrD(R421C) dimers unwind DNA with greater processivity than no-covalent wild type UvrD dimers. **(A)** Single-round stopped-flow fluorescence time courses showing DNA unwinding by crosslinked FY- UvrD(R421C) dimers of DNA substrates with different duplex DNA lengths (L) as depicted in Buffer T20-20, 500 μM ATP, 1 mM MgCl_2,_ 1 μM of DNA hairpin trap, 100 nM dimer, 25 nM DNA (post-mix) at 25°C. Analysis was performed by fitting the lag times vs. L, to a straight line (unwinding rate (bp/sec) = 1/slope). **(B)** Maximal DNA unwinding amplitudes from panel **A** plotted vs L and fit to Eq. 4 to obtain the average number of base pairs unwound, <N>. **(C)** Time courses for DNA unwinding of 18 bp and 50 bp duplexes by crosslinked FY-UvrD(R421C) dimers (blue & black) vs. wild type *Ec* UvrD (red) at excess enzyme to DNA ratios of 4:1 and 5:1 (crosslinked dimers per DNA) (100 nM crosslinked dimers, 25 nM DNA) or 8:1 (monomers per DNA, 200 nM wt *Ec* UvrD monomers, 25 nM DNA) as indicated. **(D)** Plot of fraction of DNA molecules unwound as a function of FY-UvrD(R421C) dimers (orange- no DTT) or FY-UvrD(R421C) monomers (blue- 1 mM DTT) at constant 25 nM DNA. The black curve shows the best non-linear least fit of the dimer data (no DTT) to a one-to-one binding isotherm. **(E)** DNA unwinding time courses showing that crosslinked FY-UvrD(R421C) dimers unwind DNA under conditions of excess of DNA (25 nM) over dimers (6.25 nM) (no DTT), whereas no DNA unwinding is observed under the same conditions for FY-UvrD(R421C) monomers (1 mM DTT).

**Figure 4A** shows the time courses for the stopped-flow DNA unwinding experiments with fully crosslinked FY-UvrD(R421C) dimers. A DNA unwinding rate of 119 ± 2 bp/s (Table 2) was determined from the inverse slope of a plot of the lag-time vs. dsDNA duplex length[8, 70]. This is slightly higher than the rate determined previously for non-covalent wild type UvrD dimers of 80 ± 30 bp/s [43, 51]. Surprisingly, crosslinked FY-UvrD(R421C) dimers initiate DNA unwinding much slower than wild type (wt) *Ec* UvrD dimers [43], taking ∼ 50 minutes to achieve maximum unwinding amplitude (see **Figure S4**). Multiple turnover experiments (**Figure S5**), performed by mixing ATP, Protein, and DNA in different combinations shows that the crosslinked FY-UvrD(R421C) dimers readily bind and rapidly unwind DNA in seconds. The basis for the slow rate of initiation under single round conditions is not due to a slow binding rate and must be due to slow steps associated with conformational changes needed for proper initiation.

**Table 2:** DNA Unwinding Rates and Processivities of UvrD(R421C), FY-UvrD(R421C), and Cy3-FY-UvrD(R421C)

| Construct Name | Rate<br>(bps/sec) | <N>, average<br>number of bps<br>unwound | Experiment | [ATP]<br>μM |
| --- | --- | --- | --- | --- |
| FY-UvrD(R421C) | 119±2 | 163±22 | Stopped-Flow | 500 |
| UvrD(R421C) | 84±25 | 624±27 | Optical Tweezers | 50 |
| UvrD(R421C) | 148±36 | 957±39 | Optical Tweezers | 500 |
| Cy3-FY-UvrD(R421C) | 146±45 | 1199±37 | Optical Tweezers | 500 |

**Figure 4B** shows the maximum DNA unwinding amplitudes from **Figure 4A** plotted as a function of DNA duplex length. The data in **Figure 4B** were fit to Eq. (4) to obtain <N>, the average number of base pairs unwound per binding event (Table 2), which is a measure of DNA unwinding processivity. The value of <N> = 163 ± 22 bp (Table 2), is ∼ten-fold higher than measured previously for non-covalent wt *Ec* UvrD dimers (<N>=14±3 bp)[71]. **Figure 4C** shows this directly by comparing the unwinding time courses for DNA substrates with 18 and 50 bp duplexes (3’-(dT)_20_-18 bp, 3’-(dT)_20_-50 bp) for both wt *Ec* UvrD dimers and crosslinked FY-UvrD(R421C) dimers under conditions such that the DNA substrates are saturated with enzyme. The crosslinked FY-UvrD(R421C) dimers unwind nearly 100% of the 18 bp DNA substrate whereas the non-covalent wt *Ec* UvrD dimers unwind only about 50% of the 18 bp DNA substrate. Similarly, the crosslinked FY-UvrD(R421C) dimers can unwind ∼78% of the 50 bp DNA substrate, whereas the wt *Ec* UvrD dimers show no unwinding of the 50 bp DNA substrate. Hence crosslinked UvrD dimers have much higher DNA unwinding processivity than wt UvrD dimers.

We also measured the DNA unwinding amplitudes as a function of the ratio of protein to DNA substrate (3’-(dT)_20_-18bp) at a constant DNA concentration of 25nM for fully crosslinked FY- UvrD(R421C) dimers (orange-no DTT) (**Figure S6A)** vs. un-crosslinked FY-UvrD(R421C) (blue – plus DTT) (**Figure S6B**) and these are plotted in **Figure 4D**. The data for the crosslinked FY-UvrD(R421C) dimers are well described by a one-to-one binding model with equilibrium association constant, K = 2.2 ± 0.8×10^7^ M^-1^. Significantly, **Figure 4E** shows that the crosslinked FY-UvrD(R421C) dimers (no DTT) retain helicase activity under conditions of excess DNA (4 DNA/dimer), whereas no DNA unwinding is observed by monomers (plus DTT). The loss of helicase activity for un-crosslinked enzyme results from the fact that under conditions of excess DNA, non-covalent dimers of wt UvrD [43, 44], wt Rep [27] and wt *Bst* PcrA [25] will dissociate and bind DNA as monomers that have no helicase activity. The small extent of DNA unwinding observed in the presence of DTT (**Figure 4D** and **Figure S6B**) is due to the fact that FY-UvrD(R421C) monomers retain some ability to dimerize at high concentrations. However, UvrD(R421C) shows a lower ability to dimerize than wt UvrD indicating that the R421C substitution destabilizes non-covalent dimerization. **Figure S7** shows that wt *Ec* UvrD dimers are more stable than UvrD(R421C) monomers in both the presence and absence of reducing agent.

### A single crosslinked UvrD(R421C) dimer can unwind thousands of DNA base pairs

We next examined the DNA unwinding processivity of unlabeled crosslinked UvrD(R421C) dimers in more detail by performing single DNA molecule optical tweezer experiments using a LUMICKS C-Trap. We first trapped streptavidin-coated beads within each of the two traps, and flowed in DNA in PBS buffer until a dsDNA, 17,853 bp in length, was captured via a 3x-biotin tag on the 5’- ends of both DNA strands. The DNA construct contained two nicks separated by 5,005 base pairs on one strand of the duplex DNA. The DNA was then stretched to dissociate the 5,005 nt ssDNA between the two nicks to form a ssDNA gap as depicted in **Figure 5A**. We confirmed formation of the ssDNA gap by the change in the force-extension profiles as shown in **Supplemental Figure 8**. The DNA, under a constant tension of 10 pN, was then moved into a channel of the flow-cell containing Buffer T20-20, 0.5mM ATP and 1mM MgCl_2_ and 1-3 nM of a UvrD(R421C) sample containing 45% crosslinked dimer. Any unwinding of the 6396 bp DNA was detected as movement of the mobile bead (M) away from the stationary bead (S) due to the conversion of dsDNA into ssDNA as depicted in **Figure 5A**. The number of base pairs unwound is calculated by comparing the distance moved by the mobile bead to the total distance change for complete unwinding of the 6,396 bp. The latter was determined by comparing the force-extension profiles of the initial and fully unwound DNA substrates that can be modelled by the extensible Worm-Like Chain model at 10pN of force as described [30, 31] **(see Materials and Methods and Supplemental Figure 9)**.

**Figure 5.**
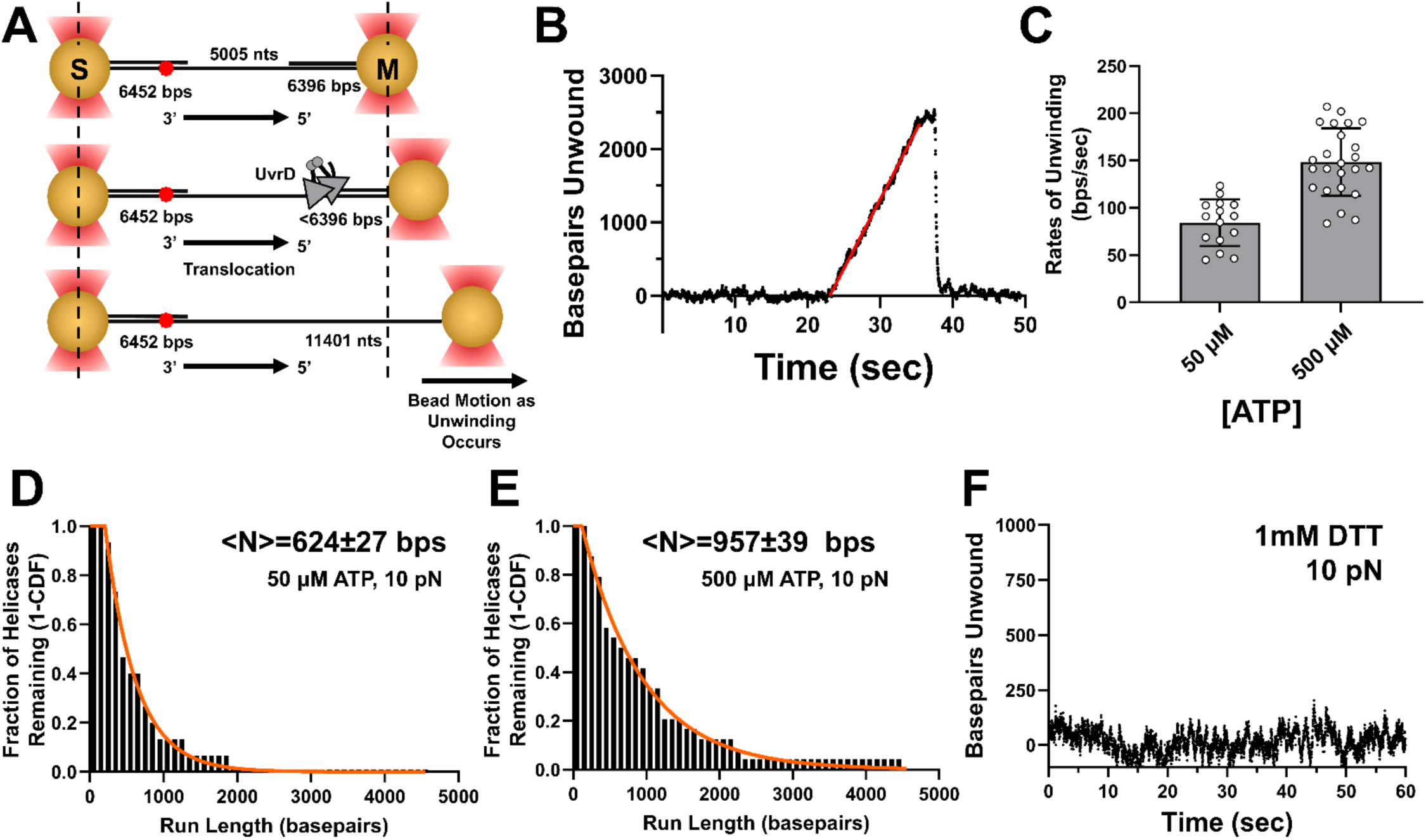
Single molecule experiments show UvrD(R421C) dimers can unwind more than one thousand base pairs. **(A)** Schematic of the LUMICKS DNA unwinding experiment using DNA with a 5,005 nt ssDNA gap flanked by two dsDNA handles, 6,452 bp and 6,396 bp in length attached to a stationary bead (S) and a mobile bead (M) while maintaining a constant force of 10pN. A fluorescent 647N Atto dye is incorporated into the 6,452 bp DNA duplex 177 bp from the 5’-ss/dsDNA junction. Cy3-FY-UvrD(R421C) binds to the ssDNA gap and translocates 3’-to-5’ toward the 3’- ss/dsDNA junction of the 6,396 DNA duplex. Upon DNA unwinding by the helicase the dsDNA is converted to ssDNA and the mobile bead will move away from the stationary bead to maintain a constant force. Experiments were performed in Buffer T20-20 plus 1mM MgCl_2_, 50µM or 500µM ATP at 25°C**. (B**) A DNA unwinding event for a crosslinked UvrD(R421C) dimer resulting from the unwinding of 2,462 base pairs at a rate of 190 bp/s (linear red line) in 500 µM ATP. **(C)** Rates of DNA unwinding by crosslinked UvrD(R421C) dimers are 84 ± 25 bp/s (n=15) at 50 µM ATP and 148 ± 36 bp/s (n=24) at 500 µM ATP (Table 2). Processivity of crosslinked UvrD(R421C) dimers at **(D)** 50µM ATP and **(E)** 500µM ATP at 10pN. The data were fit to Eq. 7 to obtain the average number of base pairs unwound, <N>. Errors are the standard error of the fit (Table 2). **(F)** Example trace in 500µM ATP showing that a UvrD(R421C) monomer formed by adding 1mM DTT is unable to unwind duplex DNA.

We observed multiple DNA unwinding events by individual crosslinked UvrD(R421C) dimers. **Figure 5B** shows one example in which a crosslinked UvrD(R421C) dimer unwinds nearly 2,500 bp. Upon dissociation of the helicase, the DNA re-hybridizes rapidly to its initial fully duplex state. No such events are observed in the absence of enzyme or ATP. A single DNA duplex can be unwound multiple times as long as the DNA reanneals unless the DNA is fully unwound resulting in release of one DNA strand. Rates of unwinding (Table 2) were determined by a linear least squares fit of the ascending portion of the data (red line in **Figure 5B**). All of the DNA unwinding events are shown in **Supplemental Figures 10** (50 μM ATP) and **11** (500 μM ATP). **Figure 5C** shows a plot of the rates of DNA unwinding from each individual event at 50 µM and 500 µM ATP, with an average unwinding rate of 84 ± 25 bp/s at 50 µM ATP (Table 2) and 148 ± 36 bp/s at 500 µM ATP (Table 2) indicating that the DNA unwinding rate is ATP dependent. Plots of the fraction of helicase dimers remaining bound to the DNA as a function of the number of base pairs unwound by each dimer (**Figure 5D** and **5E**) were fit to a single exponential decay (**Eq. 7**) to obtain the average extent of DNA unwinding, <N>, by a UvrD(421C) dimer (Table 2). Processivity increases with [ATP], with <N> = 624 ± 27 bp at 50 µM ATP (Table 2), while <N> = 957 ± 39 bp at 500 µM ATP (Table 2). Upon adding 1 mM DTT, no DNA unwinding events were observed indicating that monomers of UvrD(R421C) do not unwind DNA **(Figure 5F)** consistent with previous studies [43, 44, 55] and the ensemble studies reported here. The rates of DNA reannealing or rehybridization after DNA unwinding were determined by linear fits to the data (blue lines in **Supplemental Figures 10** and **11).** These rates of DNA rehybridization after an unwinding event were much faster than the DNA unwinding rates and independent of [ATP] (450 ± 336bp/s at 500 µM ATP vs. 605 ± 666bp/s at 50 µM ATP, p-value=0.56) (**Supplemental Figure 12,** Table 2). This suggests that reannealing is unrelated to enzyme activity.

We also performed single molecule DNA unwinding experiments using the Cy3-labeled crosslinked FY-UvrD(R421C) dimers, enabling us to detect DNA unwinding while also directly monitoring the enzyme on the DNA via its Cy3 fluorescence. **Figure 6A** (upper panel) shows kymographs for three crosslinked Cy3-FY-UvrD(R421C) dimers each of which is associated with a distinct DNA unwinding event (lower panel) in Single-Molecule Imaging Buffer (500 µM ATP). The unwound DNA fully rehybridizes after dissociation of the dimeric helicase from DNA after the first two unwinding events and partially rehybridizes after the third. The rates of DNA unwinding from 14 such events are shown in **Figure 6B**, with an average rate of 146 ± 45 bp/s (Table 2), which is the same as observed for the unlabeled crosslinked UvrD(R421C) dimers at the same [ATP] of 500 µM **(Figure 5C)**. All DNA unwinding events and associated kymographs are presented in **Figure S13**, showing the fits used to determine the rates of unwinding and DNA rehybridization. DNA rehybridization rates are 1405 ± 904 bp/s (Table 2) (see **Figure S12**) and are significantly higher than the rates of translocation of 200 ± 77 nt/s (**Figure 3B**). Taken together, the differences in our measured rates for ssDNA translocation and DNA rehybridization, along with the absence of bidirectional movement of Cy3-FY-UvrD(R421C) dimers, suggest that strand-switching of the dimers does not occur and that DNA rehybridization is not mediated by enzyme activity. **Figure 6C** shows a plot of the fraction of helicases remaining bound to DNA (1-CDF) vs. the number of base pairs unwound by each dimer. After a plateau region, this plot is well described by a single exponential decay from which we estimate that Cy3-FY-UvrD(R42C) dimers can unwind an average of <N> = 1199 ± 37 bp per unwinding event (Table 2), which is similar to <N> for the crosslinked UvrD(R421C) dimers shown in **Figure 5E**. These results suggest that there is no significant effect of the tags or Cy3 label on dimer helicase activity.

**Figure 6.**
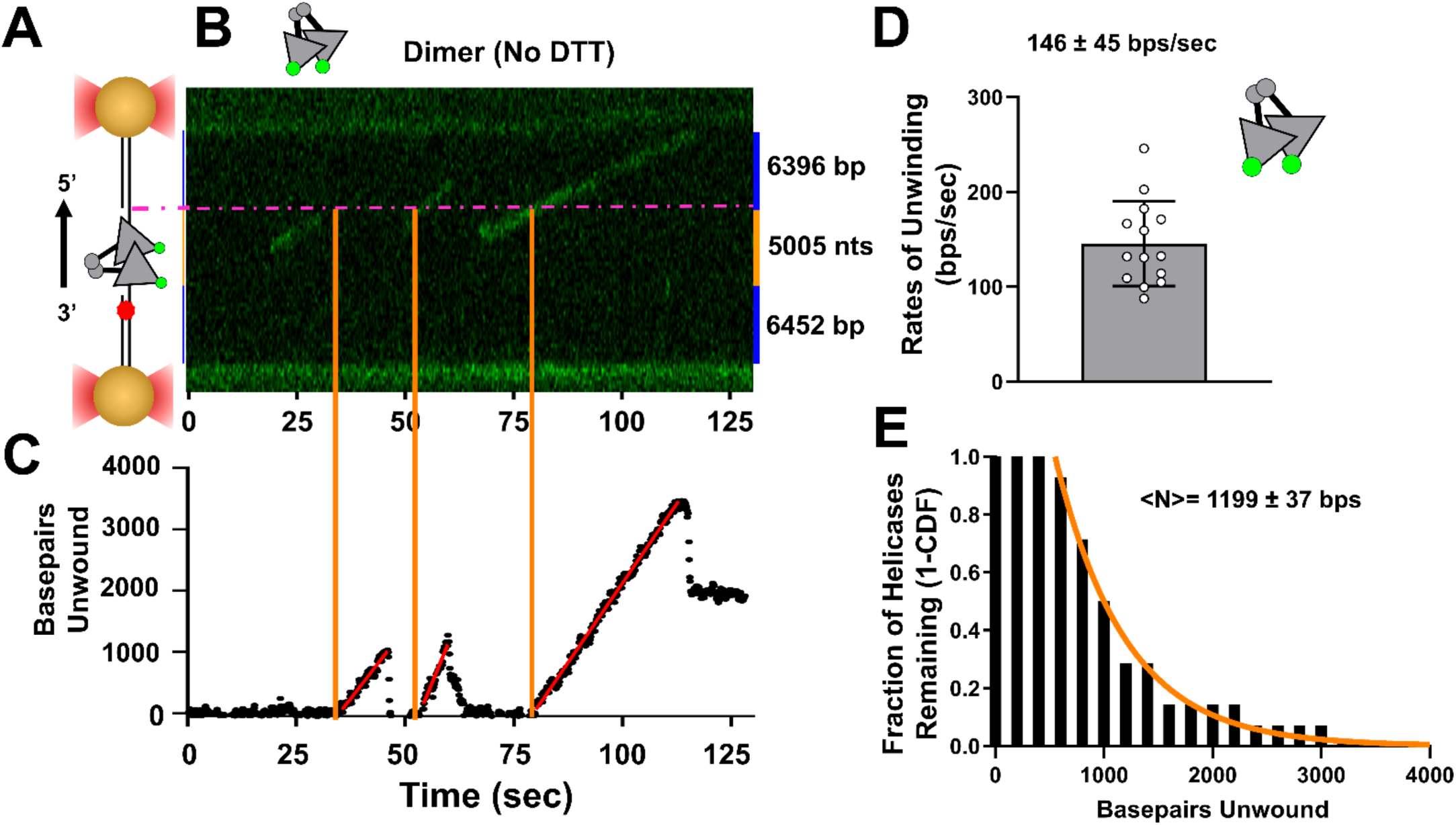
Combined fluorescence optical tweezer experiment shows crosslinked Cy3-FY-UvrD(R421C) dimers can unwind thousands of base pairs. Experiments were performed as in Figure 5A with but with Cy3-labeled crosslinked FY-UvrD(R421C) dimers and confocal imaging utilizing a 532 nm excitation in single-molecule imaging buffer (500 µM ATP). **(A)** Schematic of gapped DNA substrate. (**B**) Kymographs showing DNA unwinding by three crosslinked Cy3-FY-UvrD(R421C) dimers. The orange vertical lines indicate the positions where each dimer initiates DNA unwinding at the 3’-ss/dsDNA junction (dot-dash magenta line). (**C**) Plot of the number of base pairs unwound as a function of time and the linear fits (red lines) to determine the unwinding rates of each event. **(D)** DNA unwinding rates measured for individual crosslinked Cy3-FY-UvrD(R421C) dimers show an average rate of 146 ± 45 bp/sec (mean ± standard deviation, n =14) (Table 2). **(E)** Individual crosslinked Cy3-FY-UvrD(R421C) dimers can unwind an average of <N> = 1199 ± 37 bp (Table 2), with some unwinding more than 3,000 bp.

### Cy3-FY-UvrD(R421C) monomers do not unwind DNA but exhibit specificity for a 5’-ss/dsDNA junction

We next performed single molecule (LUMICKS C-trap) experiments with Cy3-FY-UvrD(R421C) monomers on the gapped DNA to examine DNA unwinding (**Figure 7A**). **Figure 7B** shows that Cy3- labeled FY-UvrD(R421C) monomers, formed in the presence of 1mM DTT, bind and translocate along the ssDNA gap with 3’ to 5’ directionality. However, the kymographs in **Figure 7B** show that no monomers that reach the 3’-ss/ds DNA junction proceed beyond that point, but remain at the junction until they dissociate from the DNA. Furthermore, the DNA length and force placed on the DNA remains constant (**Figure 7C**) indicating that no DNA unwinding events are associated with monomers. **Figure 7B** also shows that a significant number of monomers initiate translocation at or near the 5’-ss/ds DNA junction. This is in contrast to the crosslinked dimers that initiate randomly along the ssDNA gap (**Figure 6A**). **Figure 7D** shows a plot of the number of Cy3-FY-UvrD(R421C) monomers that initiate at the 5’- ss/dsDNA junction vs. anywhere else on the ssDNA compared with the crosslinked Cy3-FY- UvrD(R421C) dimers. A Chi-squared test shows that significantly more monomers initiate at the 5’- ss/dsDNA junction in contrast to the crosslinked dimers (p-value = 0.0003). This result is consistent with previous ensemble studies showing that wt *Ec* UvrD monomers bind with specificity to both 5’- ss/ds DNA and 3’-ss/ds DNA junctions and provide a high affinity loading site for monomeric UvrD [73]. Those same studies showed that the 2B sub-domain of *Ec* UvrD is required for this binding specificity to the 5’-ss/dsDNA junction suggesting a direct interaction of the 2B sub-domain with the duplex DNA at the 5’-ss/ds DNA junction. Since UvrD dimerization occurs via the 2B sub-domains, this prevents any 2B interaction with the DNA duplex, thus eliminating specific binding of dimers to the 5’-ss/ds DNA junction [1, 24, 55].

**Figure 7.**
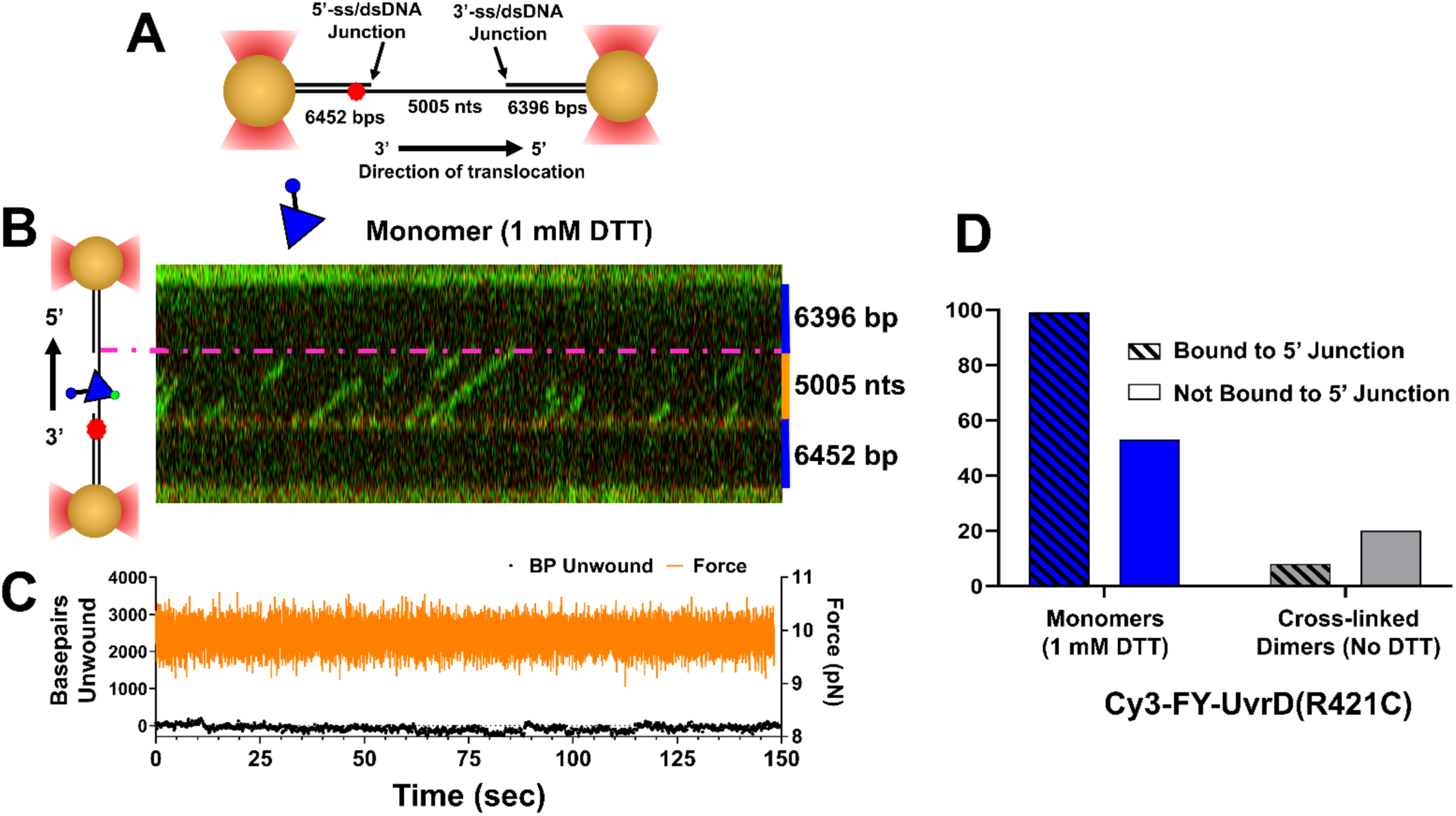
Cy3-FY-UvrD(R421C) monomers display binding specificity for 5’-ss/ds DNA junctions, do not unwind DNA. **(A)** Schematic of the ssDNA gapped DNA substrate used to examine Cy3-FY-UvrD(R421C) monomers showing the positions of the two ss/ds DNA junctions, and the 647N Atto dye near the 5’ss/ds DNA junction within the 6453 bp duplex. (**B**) Kymographs (green) show translocation of fluorescent Cy3-FY-UvrD(R421C) monomers along the ssDNA gap, but the monomers never cross the 3’-ss/ds DNA junction (magenta line) indicating that no DNA unwinding occurs (Single-Molecule Imaging Buffer plus 1 mM DTT (500 μM ATP) at 25°C). **(C)** The measured force (orange) for the experiment in panel **(B)** does not change and no base pairs are unwound (black). **(E)** Plot of the number of Cy3-FY- UvrD(R421C) monomers or crosslinked Cy3-FY-UvrD(R421C) dimers bound at the 5’-ss/ds DNA junction (near the Atto dye) or bound at some other position within the ssDNA gap. Monomers show specificity for the 5’-ss/ds DNA junction, whereas dimers show no specificity (p-value=0.0003, n= (152) monomers and (28) dimers).

## Discussion

UvrD-family monomers, such as *Ec* UvrD, *Ec* Rep, *Bst* PcrA and *Mtb* UvrD1, are rapid translocases, but lack helicase activity in the absence of force applied to the DNA [31, 32]. UvrD-family helicases can be activated by accessory proteins[50–53], by deletion of its auto-inhibitory 2B sub- domain [28, 31, 33] or by dimerization *in vitro* [27, 29, 43, 44, 46, 47]. The recently determined structure of the *Mtb* UvrD1 dimer shows that the dimeric interface of *Mtb* UvrD1 occurs between the 2B sub- domains of the two subunits and that same interface is used to form non-covalent dimers of *Ec* UvrD, *Ec* Rep and *Bst* PcrA [1, 24, 55]. Dimerization competes with an auto-inhibitory 2B sub-domain interaction with duplex DNA that occurs within the monomer and prevents DNA unwinding by UvrD-family monomers [1, 24, 55]. Placement of a Cys into the 2B sub-domain of *Ec* UvrD facilitates a redox-dependent formation of crosslinked UvrD dimers that have constitutive helicase activity [1, 55]. The same dimerization interface occurs in *Ec* Rep dimers as well as *Bst* PcrA dimers and covalently crosslinked dimers of Rep and PcrA show constitutive helicase activity with dramatically enhanced processivity [55]. This has enabled us to isolate crosslinked UvrD dimers that can be used to study both ssDNA translocation and DNA unwinding without complications from UvrD monomer activity or dimer dissociation. Here we have used these crosslinked UvrD dimers to probe the properties of individual UvrD dimers using single molecule optical tweezer methods at low protein concentrations.

The average rates of DNA unwinding measured for the crosslinked (Cy3)-FY-UvrD(R421C) dimers in the single molecule experiments (146 ± 45 bp/s) are the same within error as the ensemble rates measured using the stopped-flow (119 ± 2 bp/s) (Table 2). We find that crosslinking of UvrD dimers via its 2B sub-domain not only activates the helicase by relieving the auto-inhibitory effects of the 2B sub-domain [28, 31, 33], but also greatly enhances the processivity of DNA unwinding. On average, crosslinked UvrD dimers can unwind 1000 base pairs (<N> = 957 ± 39 bp) before dissociation, with some dimers unwinding more than 2000 base pairs. This increased processivity is presumably due to the fact that crosslinking prevents dissociation of the dimer into monomers that more readily dissociate from the DNA. In addition, monomers formed by dimer dissociation are unable to unwind DNA.

Interestingly, we measure a much higher DNA unwinding processivity in the single molecule experiments (<N> =957 ± 39 bp) than from the stopped-flow experiments (<N> = 163 ± 22 bp) at 0.5 mM ATP. Possible explanations for this difference include the fact that the trap for free protein that is included in the stopped-flow experiments may facilitate enzyme dissociation from the DNA [67]. However, there also may be multiple classes of enzymes in the population, some with lower processivity that would be detected in the stopped-flow experiments, but not in the single molecule experiments. The single molecule experiments would not be expected to detect low processivity enzymes, but only those with high processivities.

Our ability to study the same sample of UvrD that is 100% crosslinked dimer in the absence of reducing agent and 100% monomer upon addition of reducing agent has allowed us to directly compare their ssDNA translocation for the first time. The interpretation of studies of ssDNA translocation by non-covalent UvrD dimers has been difficult since monomers are always present in the population at concentrations that populate dimers [43, 44, 56] and monomers can also translocate rapidly along ssDNA [7–9, 25, 67, 68]. Furthermore, UvrD tetramers also form at higher UvrD concentrations [56]. Surprisingly we find that the average rates of ssDNA translocation of crosslinked dimers and monomers are the same, although crosslinked dimers have a much higher ssDNA translocation processivity than monomers. This raises the possibility that under the conditions of our experiments dimers translocate along ssDNA using primarily one subunit, although this hypothesis needs further testing. However, ssDNA binding to Rep dimers has been shown to exhibit negative cooperativity [35, 36, 39, 74–76], suggesting that Rep dimers might translocate on ssDNA using primarily one subunit. The simplest explanation for the higher dimer processivity is that the second subunit is always available to bind the ssDNA when the translocating subunit dissociates. Hence, the presence of the two subunits decreases the macroscopic dissociation rate from the DNA allowing the dimer to remain bound to the ssDNA for longer times. The increased processivity may also result from intra-subunit communication between the subunits during translocation, as such subunit communication is evident for both Rep and UvrD dimers [47]. Those results and others suggest a “subunit switching” mechanism for Rep translocation along ssDNA [24, 36, 74, 77].

The rates of ssDNA translocation for monomeric FY-UvrD(R421C) measured by the ensemble stopped-flow approach are the same within error as those measure in the single molecule experiments and the same as those measured for wt *Ec* UvrD monomers [9, 67, 68], indicating that the modifications used to tag and label UvrD do not affect ssDNA translocation. Surprisingly, crosslinked dimeric FY- UvrD(R421C) displayed unanticipated ssDNA translocation time courses in the stopped-flow experiments. In contrast to the monomer translocation time courses, which indicated random initiation on the (dT)_L_ ssDNA, the translocation time courses for the dimers displayed an acute peak that moved to longer times as the (dT)_L_ length increased. This suggests that some fraction of the crosslinked FY- UvrD(R421C) dimers preferentially initiate at a unique site on the ssDNA, likely the 3’-end. The single molecule studies also did not show evidence of pausing and/or major changes in ssDNA translocation rates for either monomeric or dimeric FY-UvrD(R421C). Notably, we also did not observe effects of force on ssDNA translocation for either monomeric or crosslinked FY-UvrD(R421C) dimers, **(Figure 3B** and **C)**, consistent with previous single-molecule studies on monomeric UvrD that also observed force-independent ssDNA translocation rates [45, 78].

The single molecule experiments show that the monomeric forms of our UvrD variant are unable to unwind DNA, consistent with previous studies [43, 45], but also confirmed that Cy3-FY-UvrD(R421C) monomers can bind and processively translocate along ssDNA. In addition, the experiments show directly that FY-UvrD(R421C) monomers exhibit specificity for binding to a 5’-ss/dsDNA junction. In fact, previous stopped-flow and DNA binding study indicate that UvrD monomers show specificity for binding both a 5’-ss/ds DNA junction as well as a 3’-ss/ds DNA junction [73]. The crosslinked UvrD dimers do not display this DNA junction specificity. This specificity appears to be due to the fact that the 2B-domain in the monomer is free to interact with the duplex DNA at both junctions, while the 2B sub-domain is involved in dimerization and thus prevented from interacting with the duplex DNA.

Previous single molecule studies have reported reannealing events during the course of DNA unwinding by some helicases [32, 49, 78]. These events have been hypothesized to occur due to “strand switching” in which a helicase, after unwinding some amount of DNA, will transfer to the complementary strand of ssDNA and translocate away from the ss/dsDNA junction. Such events have generally been observed for DNA unwinding by monomeric SF1 and SF2 enzymes in experiments in which tension is applied to the DNA. However, *S. cerevisiae* Pif1 helicase also exhibits these DNA reannealing events, yet strand switching has been ruled out in that case [79]. We have not observed any such “strand switching” events during DNA unwinding by the crosslinked UvrD dimers. Our single molecule experiments indicate that when DNA unwinding events end, abrupt DNA rehybridization of the DNA duplex occurs. The rate of rehybridization is much more rapid than the rate of ssDNA translocation by the UvrD dimer and is not affected by [ATP] suggesting that rehybridization does not involve enzyme activity. If such strand switching required interaction of the 2B sub-domain with the complementary ssDNA, the constrained conformation of the 2B-domains within the crosslinked dimers would prevent such transient interactions. Studies with Rep-X, a variant of *Ec* Rep that contains an internal monomeric crosslink that constrains the 2B domain into a “closed” conformation also does not exhibit strand-switching [54]. However, previous studies have found that RepΔ2B, a variant of Rep without a 2B-domain, still exhibits activities that have been attributed to strand switching [31].

Finally, the studies reported here also allow us, for the first time, to directly compare the rates of DNA unwinding to the rates of ssDNA translocation by a UvrD dimer. Such comparisons have been suggested as a means to determine whether a helicase uses an “active” or “passive” mechanism to unwind DNA. An active helicase is one that plays a direct role in destabilizing the DNA base pairs during unwinding, whereas a “passive” helicase uses its directional ssDNA translocation activity to bind to and stabilize the ssDNA that is formed transiently due to thermal fraying of the DNA duplex at the ss/ds DNA junction [2, 57]. A passive helicase is expected to exhibit an almost ∼seven-fold slower rate of DNA unwinding relative to ssDNA translocation, whereas the rates of DNA unwinding are expected to be only slightly slower than for ssDNA translocation for an active helicase [58, 80]. We find that the rate of DNA unwinding by the crosslinked dimeric FY-UvrD(R421C) (146 bp/s) is only ∼20% slower than the rate of ssDNA translocation by the same protein (180-187 nt/s) indicating that the UvrD dimer functions as an “active”, rather than a “passive” helicase, playing a direct role in destabilizing the DNA duplex[2, 57, 58, 80].

## Materials and Methods

### Buffers

Buffers were prepared with reagent-grade chemicals using distilled water, further deionized with a Milli-Q purification system (Millipore Corp., Bedford, MA), and filtered through 0.22-μm filters (Millipore, Cat. # MPGP002A1)**. Buffer Tx-y** is 10 mM Tris, pH 8.3 at 25°C (unless noted), where x denotes [NaCl] (Sigma-Aldrich, Cat.# S9888) and y denotes % (v/v) glycerol (ThermoScientific, Cat.# 032450.K7) (e.g., Buffer T20-20 contains 20mM NaCl and 20% (v/v) glycerol). **Buffer Px-y** is 50 mM Tris (pH 8.3 at 25°C, unless noted) and 1 mM Na_2_-EDTA (Fisher, Cat.# S-311), where x denotes [NaCl] and y denotes % (v/v) glycerol. **Buffer R** is 20 mM Tris (pH7.4 at 22°C), 100 mM NaCl. **Single Molecule Imaging Buffer** is Buffer T20-20 plus 1 mM MgCl_2_ (Sigma-Aldrich, Cat.# M0250), 500 µM ATP (Sigma-Aldrich, Cat.# A2383), 3 mM Trolox (Sigma-Aldrich, Cat.# 238813), 0.8% (w/v) D-glucose (Sigma-Aldrich, Cat.# G8270), Glucose Oxidase (Sigma-Aldrich, Cat.# G2133; 1 mg/mL (100 units/mL), final concentration) and Catalase (Sigma-Aldrich, Cat.# G2133; (0.2mg/mL (590 units/mL), final concentration). **PBS Buffer** (Corning, Cat.# 46-013-CM) is 137 mM NaCl, 2.7 mM KCl, 1.19 mM Phosphates, 500 µM EDTA and 5 mM NaN_3_. MgCl_2_ concentrations were determined by refractive index of a stock solution in water using a refractometer (Mark II Leica Inc., Buffalo, NY) at 20°C. Stock Trolox concentrations were determined by absorbance (A_290nm_) (ε_290_ = 2,350 M^-1^ cm^-1^). ATP stocks were prepared as described[67] (ε_259_ = 1.54×10^4^ M^-1^ cm^-1^).

### Protein Purification

UvrD proteins were purified as described [13, 37] with modifications to increase solubility of the crosslinked dimers. The *Ec* FY-UvrD(R421C) mutation was introduced into plasmid pGG209ΔCys encoding a modified UvrD gene in which all six naturally occurring cysteines were changed to serine as described [1, 55]. A gBlock, purchased from IDT (Coralville, IA), containing an N-terminal tag was placed at the N-terminus of the expressed gene using a Gibson Assembly Master Mix Kit (New England Biolabs, Cat. # E2611S). The N-terminal tag consists of a leader sequence, a 9x-His-tag, a thrombin-tag, a SUMO-tag, a FLAG-tag, and a ybbR-tag, in that order (N-to-C). The leader sequence consists of amino acids Met-Val-Lys-Ile (N-to-C), which may aid in protein expression efficiency as described [81]. The SUMO tag increases solubility of the expressed protein and has a cleavage site that does not leave residual N-terminal tag residues before the FLAG tag of FY- UvrD(R421C) after cleavage with ULP1 SUMO protease.

Plasmid DNA encoding the FY-UvrD(R421C) with an N-terminal SUMO-tag was used to transform *E. coli* BL21(DE3)ΔUvrD cells, plated on LB agar plates (Lamda Biotech, Cat.# C121) with kanamycin (GoldBio, Cat.# K-120-100; 50µg/mL), and left at 37°C overnight. The next morning, colonies were resuspended in Luria Broth (RPI, Cat.# L24340) and used to inoculate five fernbach flasks each containing 1L of Luria Broth supplemented with kanamycin (25µg/mL, final concentration). Cells were grown at 37°C while shaking (225 rpm) to an OD_600_= 0.8-1.0 and protein expression was induced with 1 mM IPTG (GoldBio, Cat.# I2481C100) and grown overnight at 16°C. Cells were harvested by centrifugation (5,320xg at 4°C), and resuspended in Buffer P100-0 (pH 7.9 at 22°C) followed by centrifugation (5,509xg at 4°C). Cell pellets were resuspended in Buffer P200-0 with 10% (w/v) sucrose, 1 mM PMSF (Sigma-Aldrich, Cat.# P7626) and 0.2 mg/mL lysozyme on ice (Sigma-Aldrich, Cat.# L6876). Sodium deoxycholate (0.05% (w/v)) (Sigma-Aldrich, Cat.# D6750) was added to and incubated at 15°C for 20 minutes. NaCl was then added to 0.48M, followed by incubation at 4°C for 20 minutes with stirring. Samples were sonicated as described [37], and the cell lysate was cleared by centrifugation (25,000xg at 4°C). The supernatant was applied to a Nickel NTA Column (5 mL) (Marvelgent Biosciences, Cat.# 11-0224), in Buffer P250-20 (2mL/min), washed with Buffer P250-20 and then Buffer P250-20 plus 15mM imidazole (Acros, Cat.# 122020020) to remove non-specifically bound proteins. FY-UvrD(R421C) was eluted with Buffer P250-20 plus 500 mM imidazole. The eluate was collected, and ULP1 protease added to the sample 125:1 and dialyzed overnight against three changes of 1L of Buffer P500-20 (3.5kDa dialysis tubing). The dialyzed sample was diluted with Buffer P0-20 to a conductivity of 6-7.5mS/cm (75 mM NaCl) and applied to a Heparin Column (2x5mL), (Cytiva, Cat.# 17-0407-01). The column was washed with Buffer P75-20 and FY-UvrD(R421C) was eluted with a 50 mL NaCl gradient using Buffer P75-20 and Buffer P1000-20. Fractions containing FY- UvrD(R421C) were identified by denaturing SDS-PAGE, pooled and dialyzed against three changes of 1L Buffer P500-20 (25 kDa MWCO dialysis). The sample was then diluted with Buffer P0-20 to a conductivity of ∼6-7.5mS/cm (75 mM NaCl) and flowed through a dsDNA column (2 mL) by syringe. The flow-through was collected and applied to a ssDNA column (∼36 mL), washed with Buffer P75-20 and eluted with a 60 mL NaCl gradient (75 mM to 2M NaCl) using P75-20 and Buffer P2000-20. Denaturing SDS-PAGE was performed to identify fractions containing FY-UvrD(R421C) dimers and these were pooled and dialyzed against three changes of Buffer P75-35 with 2 mM Na_2_-EDTA (1L). FY- UvrD(R421C) concentration was determined by absorbance (ε_280nm_ = 1.07×10^6^ M (subunit)^-1^ cm^-1^), and aliquots were flash frozen in liquid nitrogen and stored at −80°C. Two milligrams of purified FY- UvrD(R421C) was obtained from 13g of cell paste using this protocol. The purified FY-UvrD(R421C) contained 96 ± 1% (mean ± standard deviation, n=3) crosslinked dimer.

### Fluorescent labeling of FY-UvrD(R421C) and dimer purification

Purified crosslinked FY- UvrD(R421C) dimers (∼96% dimers) were labeled with a Cy3 fluorophore as described [60, 61] by mixing 21 mM MgCl_2_, 55 µM Cy3-CoA (Sirius Fine Chemicals, Bremen, Germany; Cat.# SC1143), 9.2 µM Sfp protein, and 1.7 µM FY-UvrD (final concentrations) for 90 minutes on ice and then applying it to a Superdex 200 10/300 GL (GE, Cat.# 17-5175-01) size exclusion column equilibrated with 50 mM Tris (pH 8.3 at 25°C), 1 M NaCl and 10% (v/v) glycerol. The eluant was monitored by absorbance at 280 nm and 552 nm to identify the fractions containing only crosslinked FY-UvrD(R421C) dimers which were pooled. Cy3-FY-UvrD(R421C) dimer concentrations were determined by absorbance at 280 nm using Eq. (1), where (0.08x A_552_) corrects for the contribution of Cy3 at 280 nm and ε_280_ is the extinction coefficient (per subunit) of FY-UvrD(R421C) (ε_FY-UvrD(R421C),280nm_ = 1.07×10^5^ M^-1^ cm^-1^).

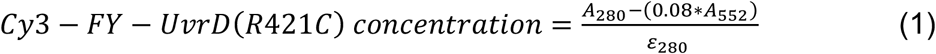

The Cy3 labeling efficiency per subunit of FY-UvrD(R421C) was calculated using Eq. (2), where the calculated FY-UvrD(R421C) (subunit) concentration, and (ε_Cy3,552_ = 1.5×10^5^ M^-1^ cm^-1^).

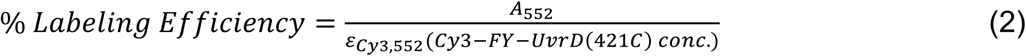

The labeling efficiency was 77±2% (mean ± std, n=3) per subunit. Assuming the two sites on FY- UvrD(R421C) are stochastically labeled, and the probability of a subunit being unlabeled is 25%, on average 93.75% of dimers have at least one Cy3 fluorophore.

### ULP1 Protein

Purification of ULP1 SUMO protease was purified as previously described. DNA plasmid, pFGET19_Ulp1 (Addgene (Cat. # 64697)), was used to recombinantly express a 6x-His tagged Ulp1 in BL21(DE3) cells (Intact Genomics, Cat.# 1051-24). ULP1 concentration was determined by absorbance (ε_280_ = 30,035 M^-1^ cm^-1^), aliquots were frozen in liquid nitrogen and stored at −80°C. Twenty-five milligrams of purified protein was obtained from 12.5g of cell paste.

### Sfp Protein

Sfp protein was purified as described[60]. Plasmid DNA, pET-Sfp (Addgene (Cat.# 159617)), was used to recombinantly express 6x-His tagged Sfp in BL21(DE3) cells. Protein concentrations were determined by absorbance (ε_280_ = 29,130 M^-1^ cm^-1^), and aliquots were frozen in liquid nitrogen and stored at −80°C. Thirty milligrams of purified protein was obtained from 5g of cell paste using this protocol.

### DNA

Oliogodeoxynucleotides used for DNA unwinding and ssDNA translocation assays were purchased from IDT (Integrated DNA Technologies; Coralville, IA). Duplex DNA stocks were prepared by annealing two complementary single-stranded (ss)DNAs in DNA Annealing Buffer and heating to 100°C in a water bath, followed by gradual cooling to room temperature overnight. ssDNA constructs used in the stopped-flow fluorescent unwinding assays are listed in Table S1 alongside the sequence and extinction coefficient used to determine the concentration by absorbance using a spectrophotometer. Extinction coefficients for the oligodeoxynucleotides are in Table S1. Oligo(dT)_L_ concentrations were determined by absorbance using ε_260nm_ = 8,100 M (nucleotide)^-1^ cm^-1^ [9, 82].

### Analytical Ultracentrifugation

Sedimentation velocity experiments were performed at 25°C at 42,000 rpm using a Proteome Lab XL-A Ultracentrifuge (Beckman Coulter, Indianapolis, IN) an An-50 Ti rotor and a double sector 12-mm Epon Charcoal-filled centerpiece. Protein samples (150-200 nM) were dialyzed vs. Buffer T20-20, and when indicated, 1mM DTT was added before loading. Absorbance data were collected at 230 nm with two replicates each scan. Data were analyzed using SEDFIT to obtain sedimentation coefficient distributions, which were converted to s_20,w_ using SEDFIT [83]. Partial specific volumes (*v̄*) for FY-UvrD at 25°C were calculated from the protein sequence using SEDFIT (*v̄* = 0.732). Buffer T20-20 density (1.0552g/mL) and viscosity (1.747 cp) was calculated using SEDNTERP [84].

### Stopped-Flow ssDNA Translocation

Stopped-Flow ssDNA translocation experiments were performed in buffer T20-20 at 25°C using an SX18MV stopped-flow (Applied Photophysics Ltd., Leatherhead, UK) as described [9, 63]. FY-UvrD(R421C) (50 nM subunit or monomer concentration), was preincubated with 5’-Cy3-(dT)_L_ (100 nM concentration) in one syringe, and reactions were initiated by 1:1 mixing with Buffer T20-20 containing 0.5 mM ATP, 1 mM MgCl_2_, and 4 mg/mL heparin (final concentrations after mixing). Heparin stocks were made and quantified as described. Cy3 fluorescence was excited at 505nm and detected with a long-pass cutoff filter (>570nm). Fluorescence time courses are averages of 8-16 repeated measurements. Monomeric FY-UvrD(R421C) translocation time courses for different lengths of DNA were globally fit to Scheme 1 by non-linear least squares analysis using Eq. (3).

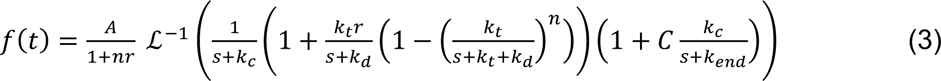

Where ℒ^−1^ is the inverse Laplace transform operator, s is the Laplace variable, A is the total protein concentration bound at time zero, r is the ratio at time zero of the protein concentration bound at any position other than the 5’-end to the protein at the 5’-end, C=(f*_end_/f_end_) is the ratio of fluorescence signals in states I_end_ and I*_end_, m is the translocation step size, and k_t_ is the translocation step rate.

### Stopped-Flow DNA Unwinding

Stopped-flow DNA unwinding experiments were performed in buffer T20-20 at 25°C and analyzed as described [51, 55, 72, 85]. FY-UvrD(R421C) was preincubated with a DNA substrate (3’-(dT)_20_ with an 18, 25, 40, or 50 bp duplex DNA) for four hours unless otherwise noted. The 5’ end of the translocating strand contains a Cy5 fluorophore while the 3’ end of the complementary strand contains a black hole quencher (BHQ2)[51]. Reactions were initiated by 1:1 mixing with Buffer T20-20 plus 0.5 mM ATP, 1 mM MgCl_2_, and 1 μM DNA trap (final concentrations after mixing). The DNA trap contained a 10 bp hairpin and a (dT)_40_ tail (5’- GCCTCGCTGCTTTTTGCAGCGAGGC-(dT)_40_-3’). Cy5 fluorescence was excited at 625 nm and detected with a long-pass filter (>665nm cutoff). Fluorescence time courses were averages of 8-18 repeated measurements. The average number of base pairs unwound per binding event, <N_bp_> was obtained from a fit of A_T_ as a function of duplex length, L, using Eq. (4) [2, 71, 72].

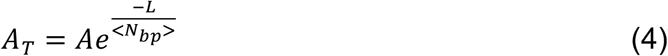

### Combined Optical Tweezer and Confocal Scanning

Single molecule DNA unwinding and translocation experiments were performed with a LUMICKS C-trap controlled with Bluelake™ (v2.5) software. The combined optical tweezer and confocal scanning microscope is outfitted with a μ-Flux™ Microfluidics System (LUMICKS) and a flow cell containing 5 channels (C1, LUMICKS). The flow cell was passivated before performing experiments by flowing PBS buffer (0.5mL at 1.6bar) through the syringes, lines, and flow cell, followed by 0.1% (w/v) BSA (Sigma, Cat.# A-6793; 0.5mL at 1.6 bar).

The experimental outline is depicted in Supplemental Figure 2. Streptavidin coated beads (4.38 μm, Spherotech Inc., Cat. # SVP-40-5) in channel 1 were captured in each of the two optical trapping lasers (1,064nm). The beads were then moved to channel 2 under flow to trap DNA biotinylated on each end to form a “dumbbell” configuration. The DNA is then moved to channel 3, which contains PBS, and force was applied to create either fully ssDNA or ssDNA gapped DNA as appropriate. Translocation experiments used a 20,425-bp DNA (LUMICKS, SKU00014). DNA unwinding experiments used a 17,853-bp tether (LUMICKS, SKU00027) that contains two nicks on the same strand, enabling the formation of a 5,005-nt ssDNA gap flanked by dsDNA handles. The dsDNA handle that is unwound in the experiments is 6,396-bp in length, and the other dsDNA handle is 6,452-bp and contains an Atto 647N fluorophore on the translocating strand within the duplex DNA 177 bp away from the ssDNA gap. The ssDNA gapped DNA was formed under flow (0.10-0.15 bar) while moving the mobile bead 9 μm away from the stationary bead. **Supplemental Figure 8** shows the change in force-extension curves upon creation of the gap.

Cumulative Distribution Function (CDF) plots of single-molecule data collected with the LUMICKS C-trap **(Figures 3E & F, 5D & E, 6E, and S3)** were constructed and fit using either Equations 5 or 7 to determine processivities. CDF plots were calculated using probabilities that the helicases will unwind or translocate by taking the binned number of observations, commonly called counts, divided by the total number of observations and integrating the data from shorter to longer lengths traveled. Probabilities are expressed as 1-CDF, to show that the probability that an enzyme will travel a specific length decreases for longer DNA lengths.

### Single Molecule ssDNA Translocation

Single molecule translocation experiments were performed as described [86]. The fully ssDNA (20,425-nt) was incubated in channel 4 containing Single Molecule Imaging Buffer, prepared by adding glucose oxidase and catalase to the buffer just before filtering the combined solution with a 0.22µm filter (GenClone, Cat No. 25-243). After filtering, FY-UvrD(R421C) protein (1-5nM) was added to the solution before loading into the microfluidics system. Kymographs were acquired using a 532 nm laser at 10% power, 27.2ms, a pixel size of 100nm while the ssDNA was maintained at a constant tension of 10 or 20 pN. Analysis was performed using Pylake(v1.6.1, LUMICKS) and Python scripts (v3.10.5) executed with Jupyter Notebooks. Individual molecule translocation trajectories were obtained by implementing the greedy tracking algorithm, which finds pixels with intensities above a threshold and subsequently refines an area of interest to determine a subpixel position before linking positions on the same trajectory together to follow the position of the molecule over time. To enable more robust tracking, the time along the kymograph image was binned by two. The average number of nucleotides translocated per binding event, <N>, was obtained by fitting the cumulative distribution function of the fraction of the remaining translocases, Y, as a function of ssDNA length travelled, x, to Eq. 5.

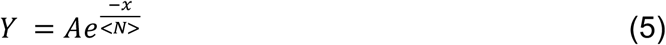

### Single Molecule DNA Unwinding

The 5,005-nt ssDNA gapped DNA substrate was incubated in channel 4 containing Single Molecule Imaging Buffer for experiments using Cy3-Labeled protein or Buffer T20-20 with 1 mM MgCl_2_, and ATP for UvrD(R421C). The UvrD(R421C) protein samples contain only ∼45% crosslinked dimer. Kymographs were acquired using both 532 nm (4-6% power) and 638 nm (2-4% power) lasers, 27.2 ms, a pixel size of 100 nm, with the ssDNA maintained at a constant tension of 10pN. Bead distance was measured while implementing a force clamp (Kp (proportional gain), 10nm/pN; Ki, (integral gain), 0.00 (nm/pN); and Kd (derivative gain), 0.0 (nm/pN) x s; Force Feedback Frequency, 31.3Hz). The movement of the bead was converted to base pairs unwound by comparing the distance the bead moved to the distance between the extensible Worm-Like Chain (eWLC) models of the initial tether and the fully unwound tether, as described [30, 31], using eq. (6) [87], where k_B_ is the Boltzmann constant and T is absolute temperature

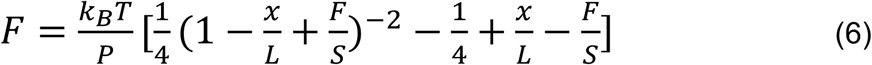

(k_B_T = 4.11 pN(nm)). P is the persistence length, P_ds_= 53nm and P_ss_= 1.2nm. S is the stretch modulus, S_ds_= 1,100pN and S_ss_= 1,000pN. x is the length of the tether (nm). The contour length (L) of the DNA tether, was calculated by multiplying h by the number of base pairs or nucleotides, where h_ds_= 0.34nm/bp and h_ss_= 0.59nm/nt. ssDNA and dsDNA eWLC simulations were performed separately and summed to recapitulate the properties of the DNA tether. The average number of base pairs unwound per unwinding event, <N>, was obtained using Eq. (7), where N1 is the offset to adjust for the plateau seen at shorter distances, and N2, determined a fit of the cumulative distribution function of the fraction of the remaining helicases, Y, as a function of the lengths of DNA unwound in base pairs, x.

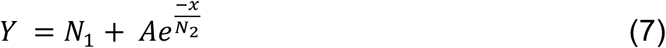

## Acknowledgements

This work was supported by NIH Grant R35 GM136632 (T.M.L). The LUMICKS C-Trap G2 was purchased with support from NIH Instrumentation grant S10OD030315. We thank Drs. Eric Galburt, Robert Galletto, Eric Tomko, and Ankita Chadda for fruitful discussions and insight.

## Author Contributions

K.N.M, B.N., A.K., and T.M.L designed research; K.N.M, B.N., A.K., and T.M.L. performed research; K.N.M, B.N., A.K., and T.M.L. contributed new reagents/analytic tools; K.N.M, B.N., A.K., and T.M.L analyzed data; K.N.M. and T.M.L wrote the paper.

## Competing Interests

The authors declare no competing interests.

## Supplemental Information

**Table S1:**
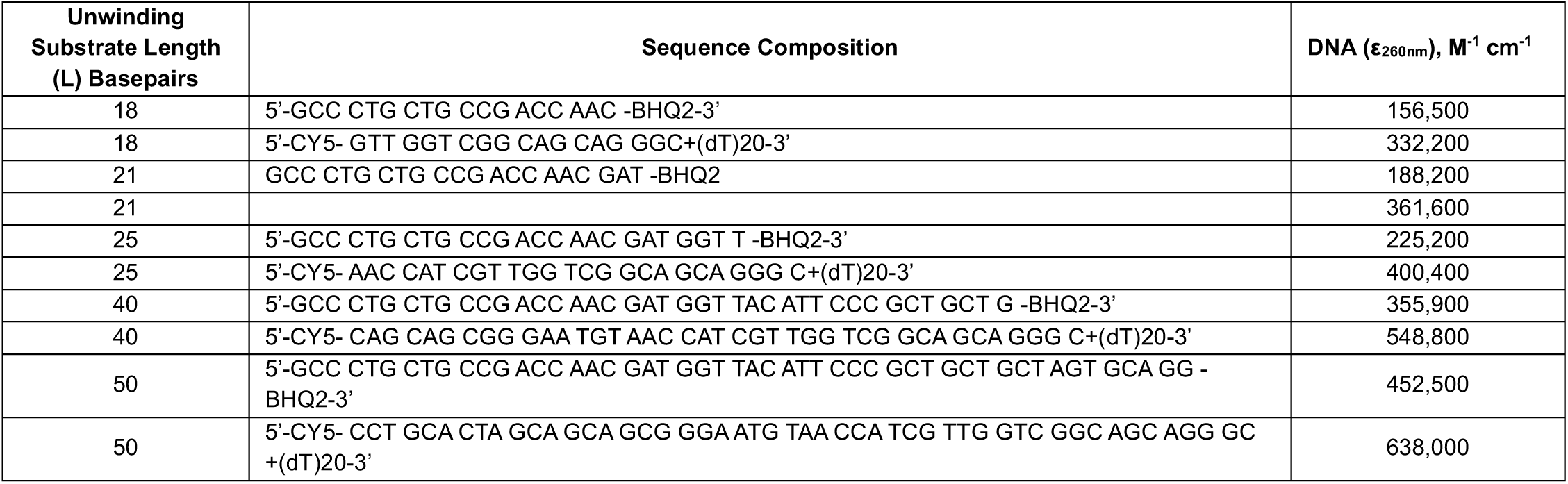
Unwinding Assay ssDNA oligomers and their extinction coefficients.

**Figure S1.**
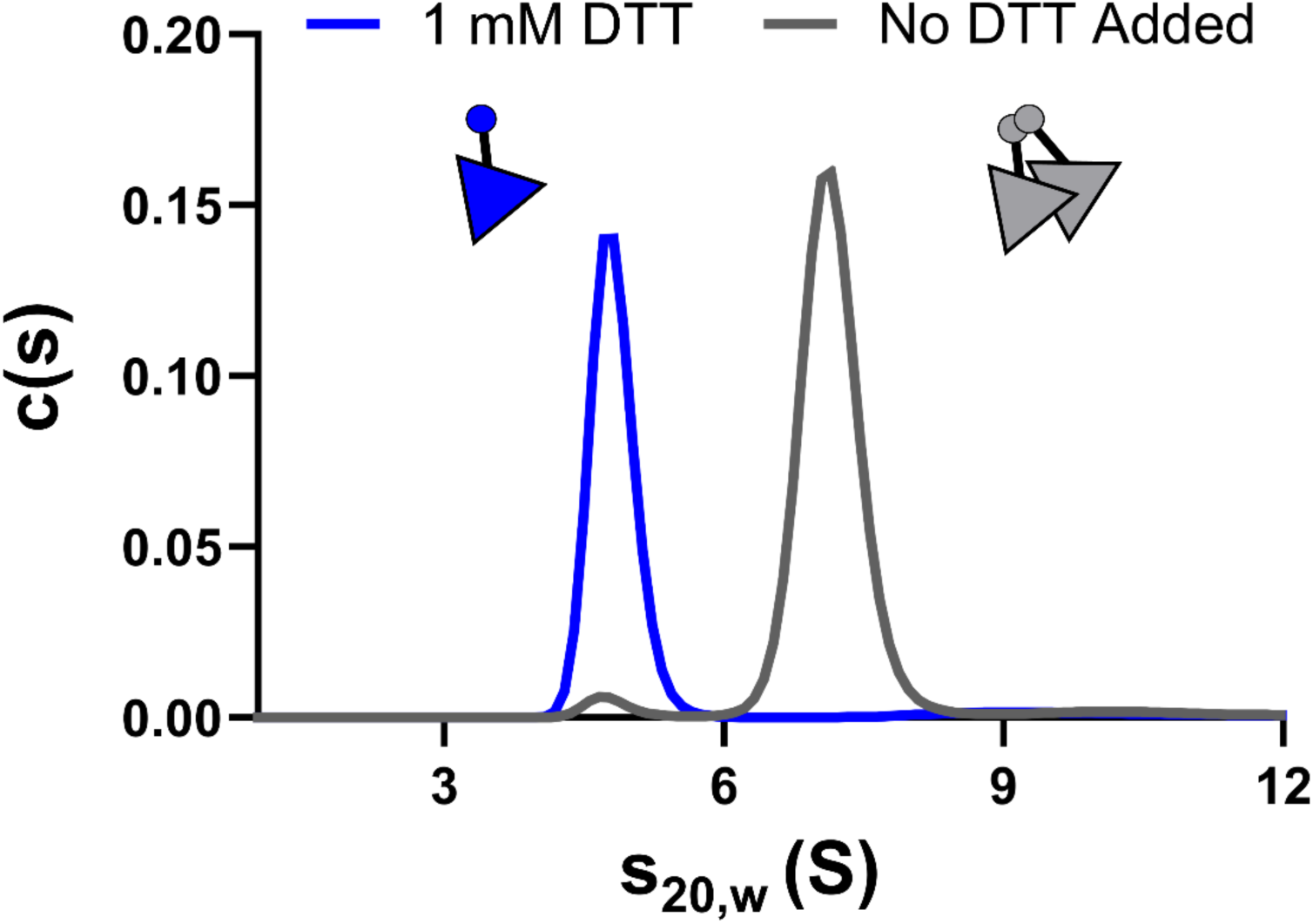
Redox-dependent dimerization of FY-UvrD(R421C). Sedimentation velocity experiments (150-200nM protein) showing FY-UvrD(R421C) dimerization is redox-dependent. FY-UvrD(R421C) is > 96% dimeric (gray) in Buffer T20-20 at 25°C (s_20,w_=7.1 S), whereas after addition of 1mM DTT the protein is entirely monomeric (s_20,w_=4.6 S).

**Figure S2.**
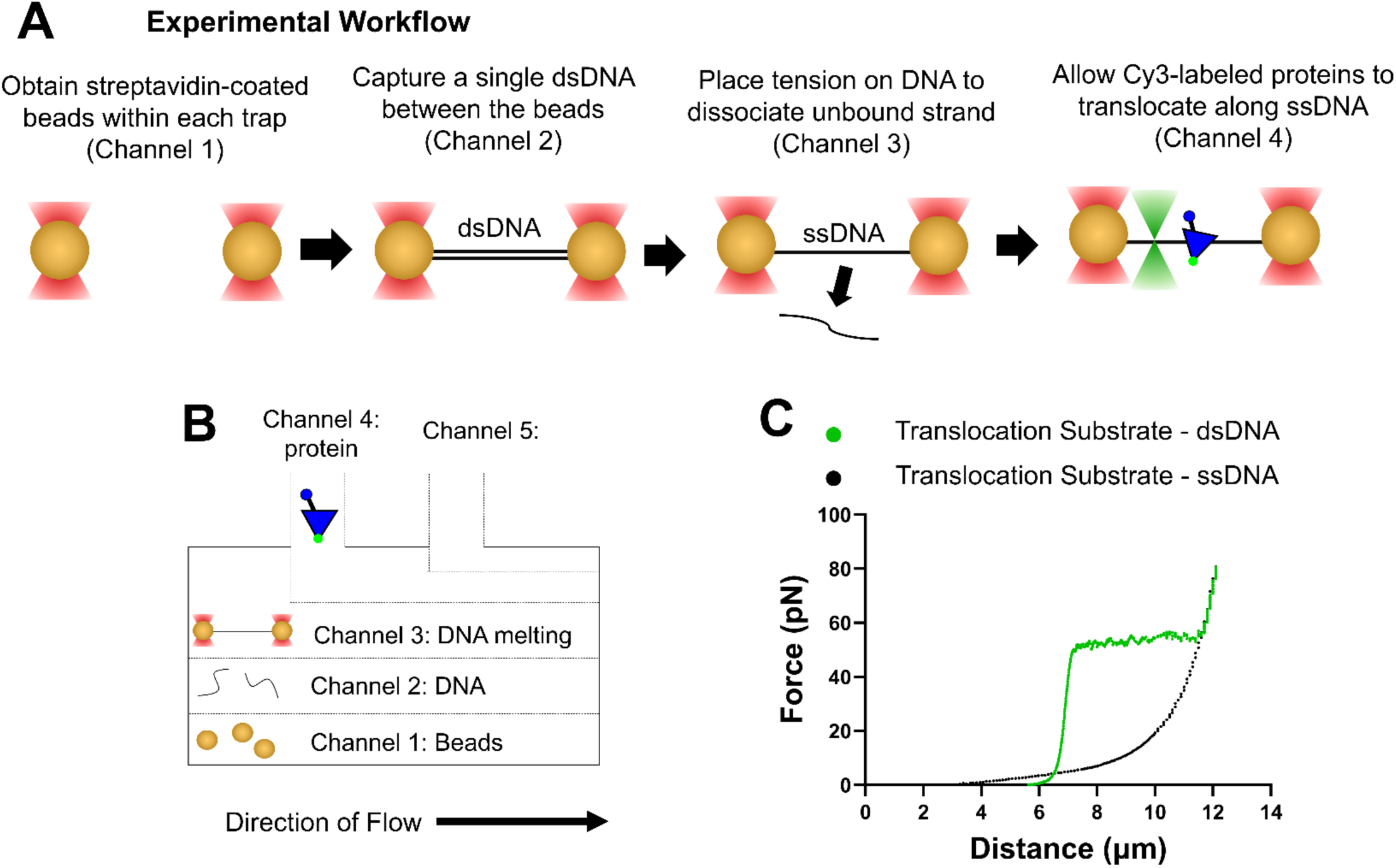
Single-molecule LUMICKS C-trap experimental set-up. **(A)** Work-flow to create a fully ssDNA begins in Channel 1 by obtaining a single streptavidin-coated polystyrene bead within each trap (see panel B). The trapped beads are moved into Channel 2 to capture a single dsDNA molecule between the beads. The dsDNA tether is moved into Channel 3 where tension is placed on the DNA until the complementary strand of DNA dissociates, producing a fully ssDNA. The ssDNA is then moved into Channel 4 for experiments. **(B)** Schematic of the LUMICKS flow cell used in the optical tweezers experiments. Channel 1 contains polystyrene beads (4.38 µm diameter) coated with streptavidin, while Channel 2 contains biotinylated dsDNA (20,425bp or 17,853bp with two nicks). In Channel 2 or 3 force is applied to the dsDNA to form a fully ssDNA or a 5,005nt ssDNA gapped DNA. Channel 4 contains 1- 3nM protein. **(C)** Force-extension curves were performed to create a fully ssDNA tether. The DNA substrate was extended while having fully dsDNA character (green circles, 20,425bp) before being relaxed (black circles) and consisting of only ssDNA (20,425nt).

**Figure S3.**
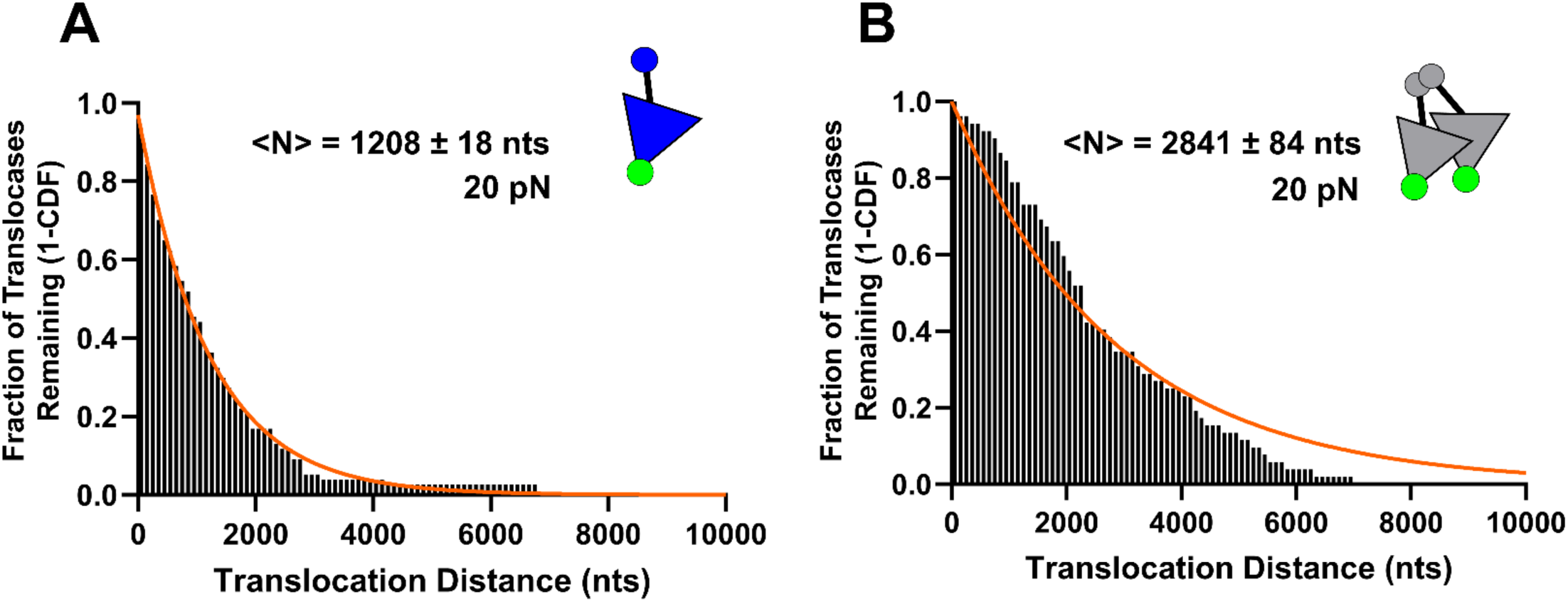
Processivities of ssDNA translocation by Cy3-FY-UvrD(R421C) monomers and dimers on ssDNA under 20pN of tension. ssDNA translocation was monitored on the 20,425nts ssDNA depicted in Figure 3A in Single Molecule Imaging Buffer at 20pN for Cy3-FY-UvrD(R421C) monomers (+ 1 mM DTT) and crosslinked Cy3-FY-UvrD(R421C) dimers (no DTT). Plots of the fraction of (**A**) monomers and (**B**) dimers remaining bound to the DNA (1-CDF) as a function of distance translocated. Data were fit with a single exponential decay Eq. (5), yielding the average number of nucleotides translocated, <N> (Table 1).

**Figure S4.**
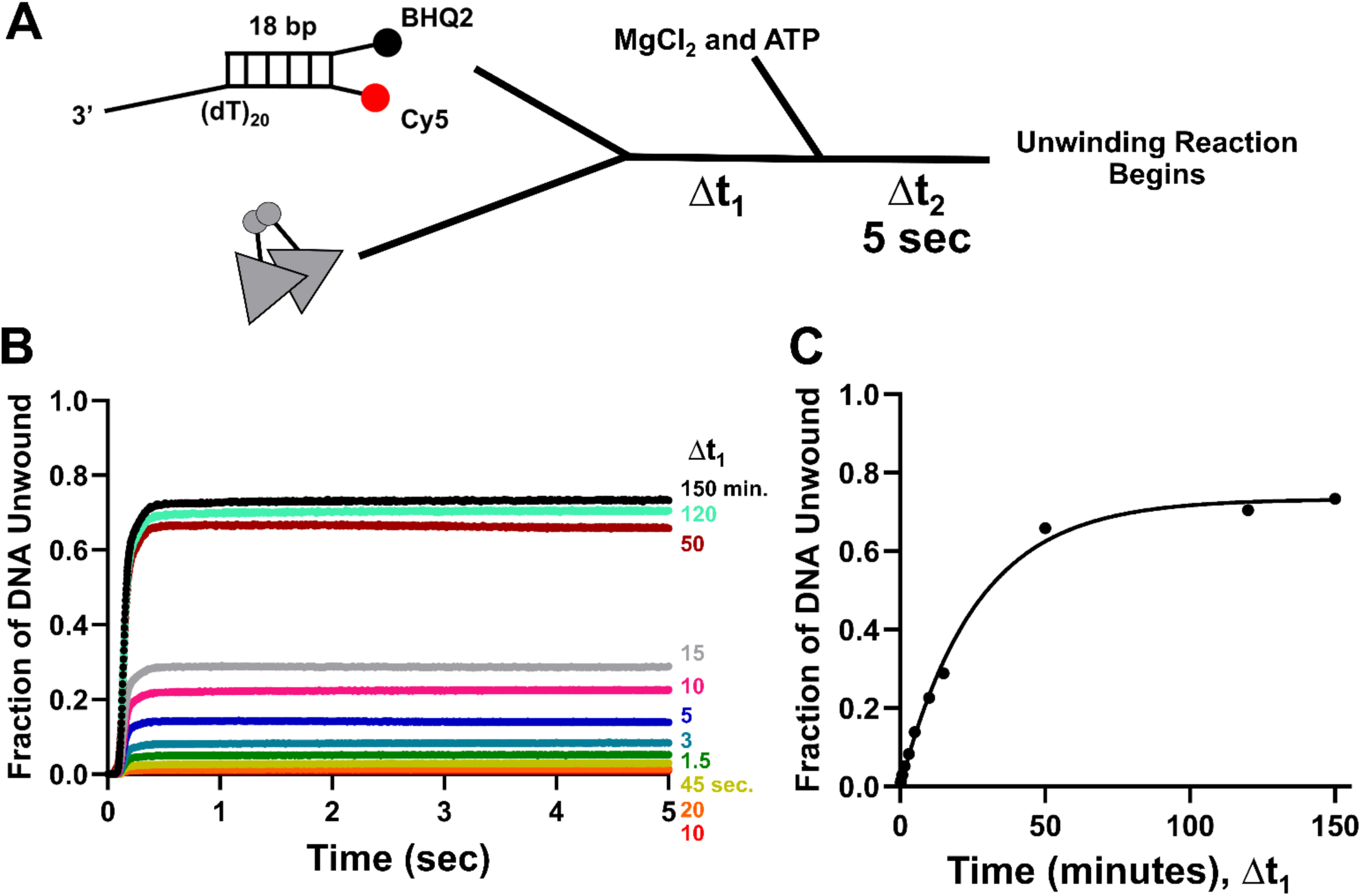
Crosslinked FY-UvrD(R421C) dimers initiate DNA unwinding slowly in single-round experiments. **(A)** Schematic of the double-mixing single-round stopped-flow experiment. In the first event, FY-UvrD(R421C) dimers (25 nM) and (dT)_20_-18bp duplex DNA (25 nM) were mixed and allowed to incubate for a variable time (Δt_1_), before mixing with 500 μM ATP, 1mM MgCl_2,_, and DNA hairpin trap (1 μM) to initiate DNA unwinding that proceeded for five seconds (Δt_2_). **(B)** DNA unwinding time courses showing the increase in DNA unwinding amplitude with increasing Δt_1_. **(C)** Plot of the maximal unwinding amplitude (filled circles) as a function of Δt_1_. The smooth curve shows the non-linear least squares fit of the data to a single exponential, y=1-e^(-Δt/^*^τ^*^)^, yielding *τ*=26±3 minutes. Experiments were performed in Buffer T20-20, 25 °C. All concentrations are post mixing.

**Figure S5.**
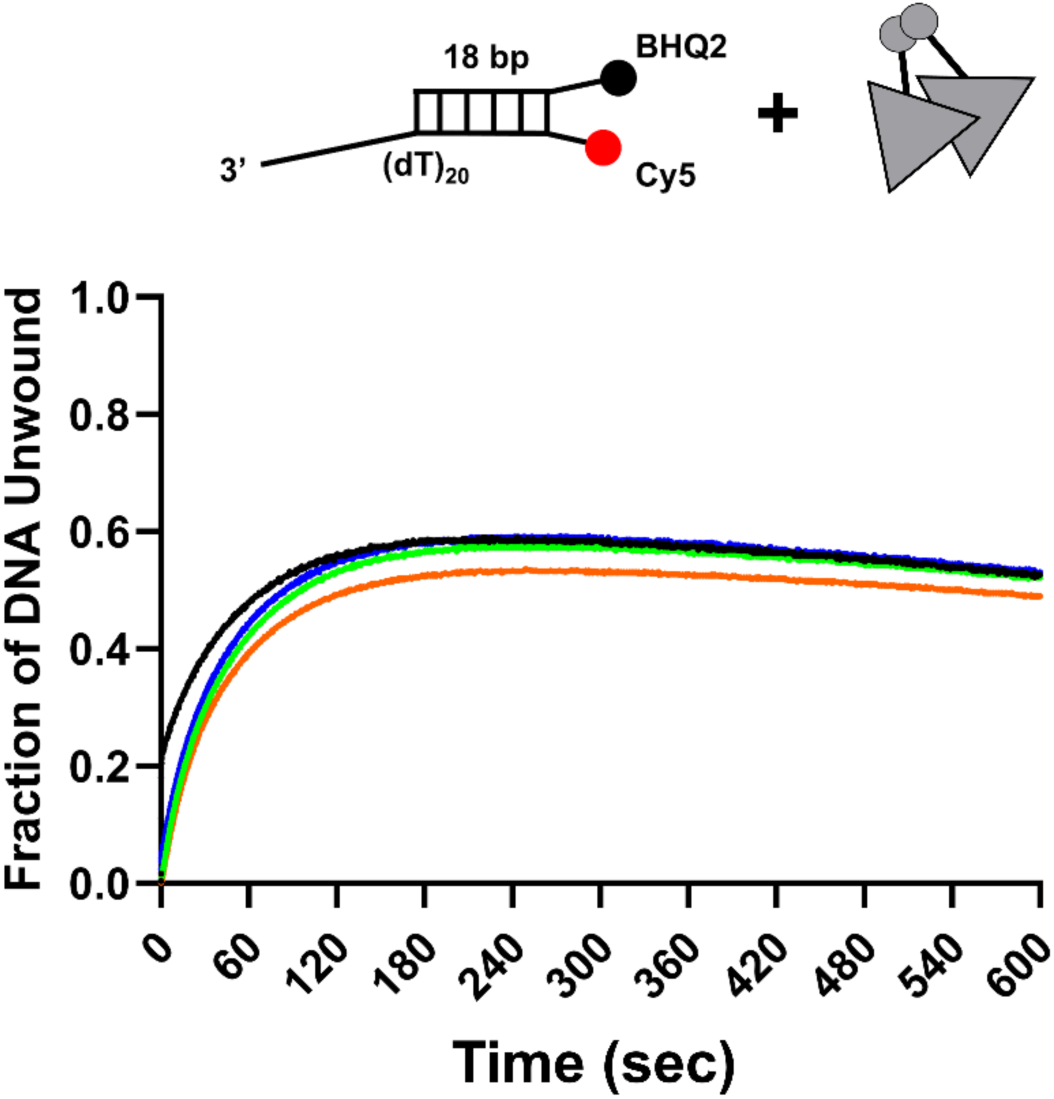
DNA unwinding by FY-UvrD(R421C) dimers in multiple-turnover experiments. Stopped-flow experiments showing (orange) mixing of (dT)_20_-18bp duplex DNA + ATP + MgCl_2_ vs. FY- UvrD(R421C) crosslinked dimers; (green) mixing (dT)_20_-18bp duplex DNA + ATP vs. FY-UvrD(R421C) crosslinked dimers + MgCl_2_; (blue) mixing ATP+MgCl_2_ vs. FY-UvrD(R421C) crosslinked dimers pre-incubated with (dT)_20_-18bp duplex DNA for two minutes; (black) mixing ATP+MgCl_2_ vs. FY- UvrD(R421C) crosslinked dimers pre-incubated with (dT)_20_-18bp duplex DNA for fifteen minutes. Experiments were performed in Buffer T20-20, 500μM ATP, 1mM MgCl_2_, 25nM DNA and 25nM FY- UvrD(R421C) [dimer] (post-mix).

**Figure S6.**
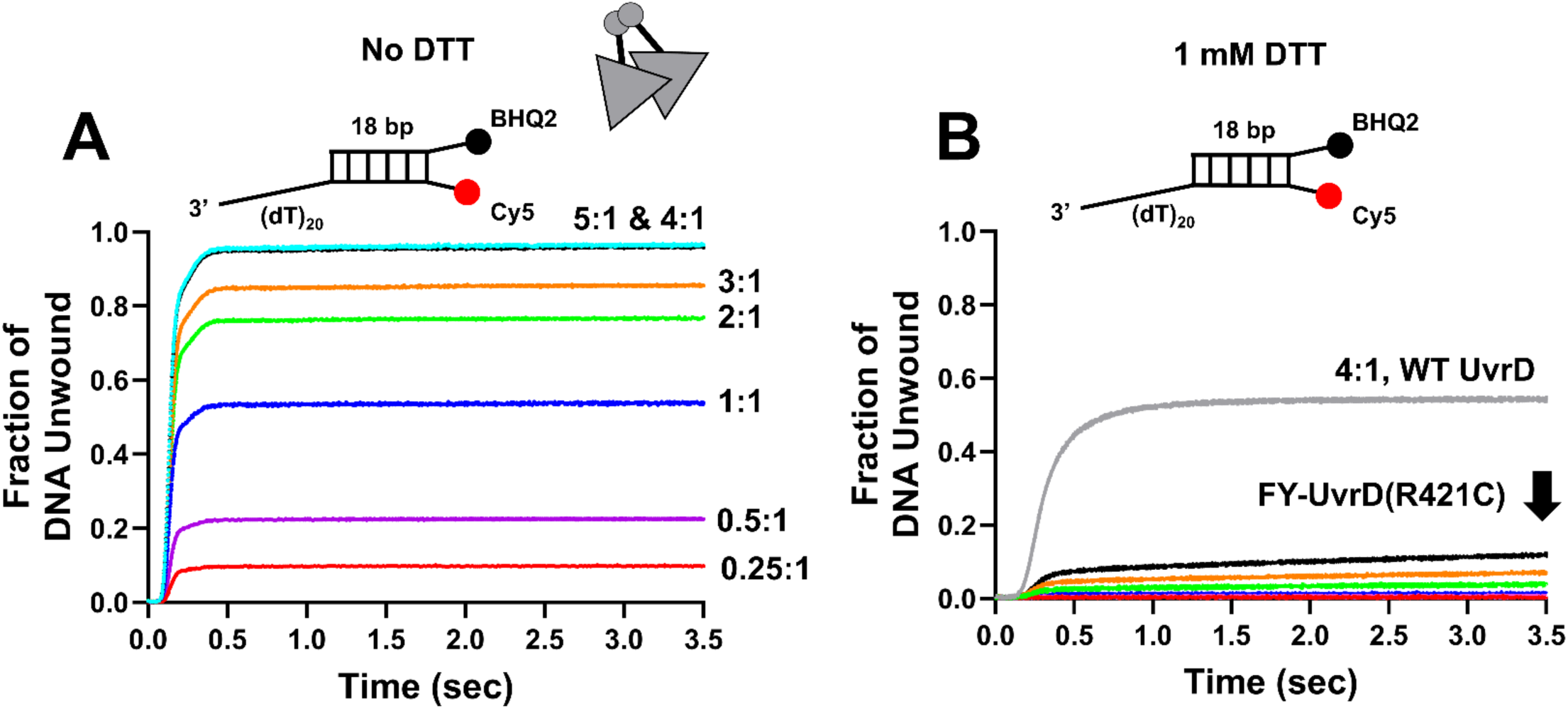
Crosslinked FY-UvrD(R421C) dimer is more processive than WT UvrD or FY- UvrD(R421C) monomers. (**A**) Stopped-flow fluorescence time courses for the data in Figure 4D showing the change in DNA unwinding amplitude as a function of **FY-UvrD(R421C) dimer concentration** at constant (dT)_20_-18bp duplex DNA (25 nM) by premixing protein and DNA for **x minutes** in Buffer T20-20, and then mixing vs. MgCl_2_+ATP+DNA trap in Buffer T20-20. **(A)** FY- UvrD(R421C) crosslinked dimers (no DTT). **(B)** FY-UvrD(R421C) (+1 mM DTT) compared with wild type *Ec* UvrD.

**Figure S7.**
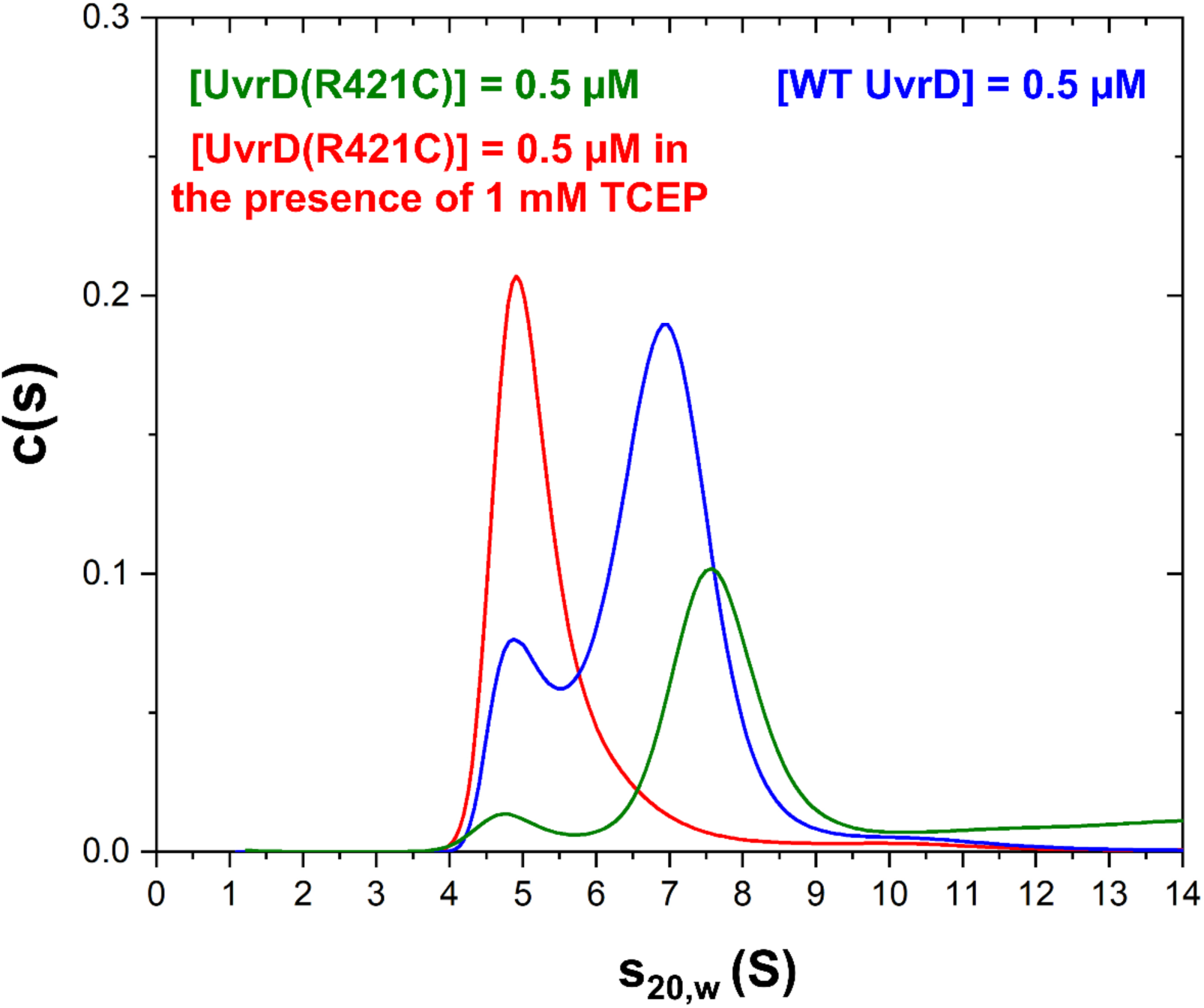
Non-covalent dimerization of FY-UvrD(R421C) is diminished compared to non-covalent wt UvrD. Sedimentation velocity experiments comparing UvrD(R421C) (green), UvrD(R421C) + 1mM TCEP (red), and wt Ec UvrD (blue) at 500 nM protein (monomer concentration) in BufferT20-20 show that UvrD(R421C) in the presence of reducing agent (1mM TCEP) has a weakened ability to dimerize compared with wt UvrD.

**Figure S8.**
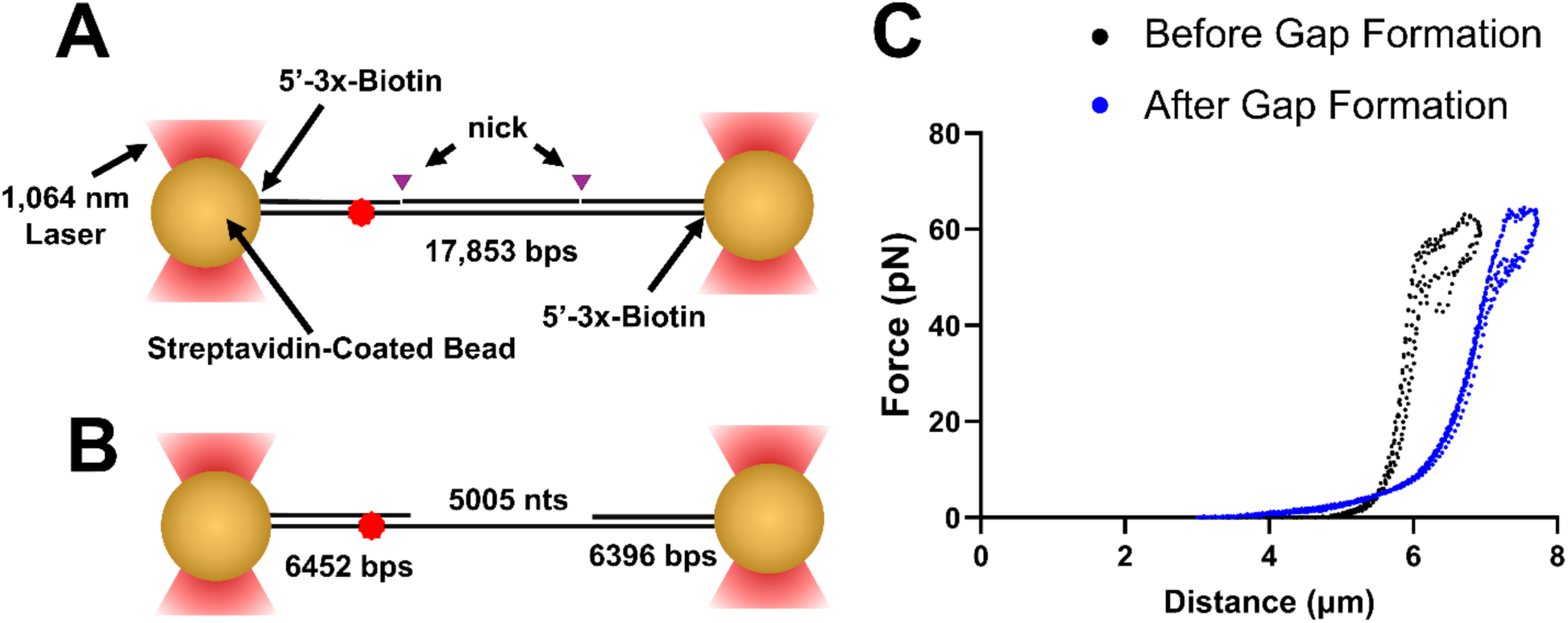
Single stranded DNA gapped formation in LUMICKS C-trap. **(A)** Schematic of the 17,853 bp doubly-nicked DNA used to form the 5,005 nt ssDNA gapped DNA. Each 5’-end of the dsDNA has a 3x-Biotin linkage to the streptavidin-coated polystyrene bead. **(B)** Upon applying force to the dsDNA the complementary DNA between the nicks dissociates, leaving a 5,005 nt ssDNA gap and two dsDNA handles, 6452 bps and 6396 bps in length. **(C)** Force-extension profiles of the (Black)- doubly nicked dsDNA used in Figures 5, 6 and 7; (Blue)- after formation of the 5,005 nt ssDNA gapped DNA.

**Figure S9:**
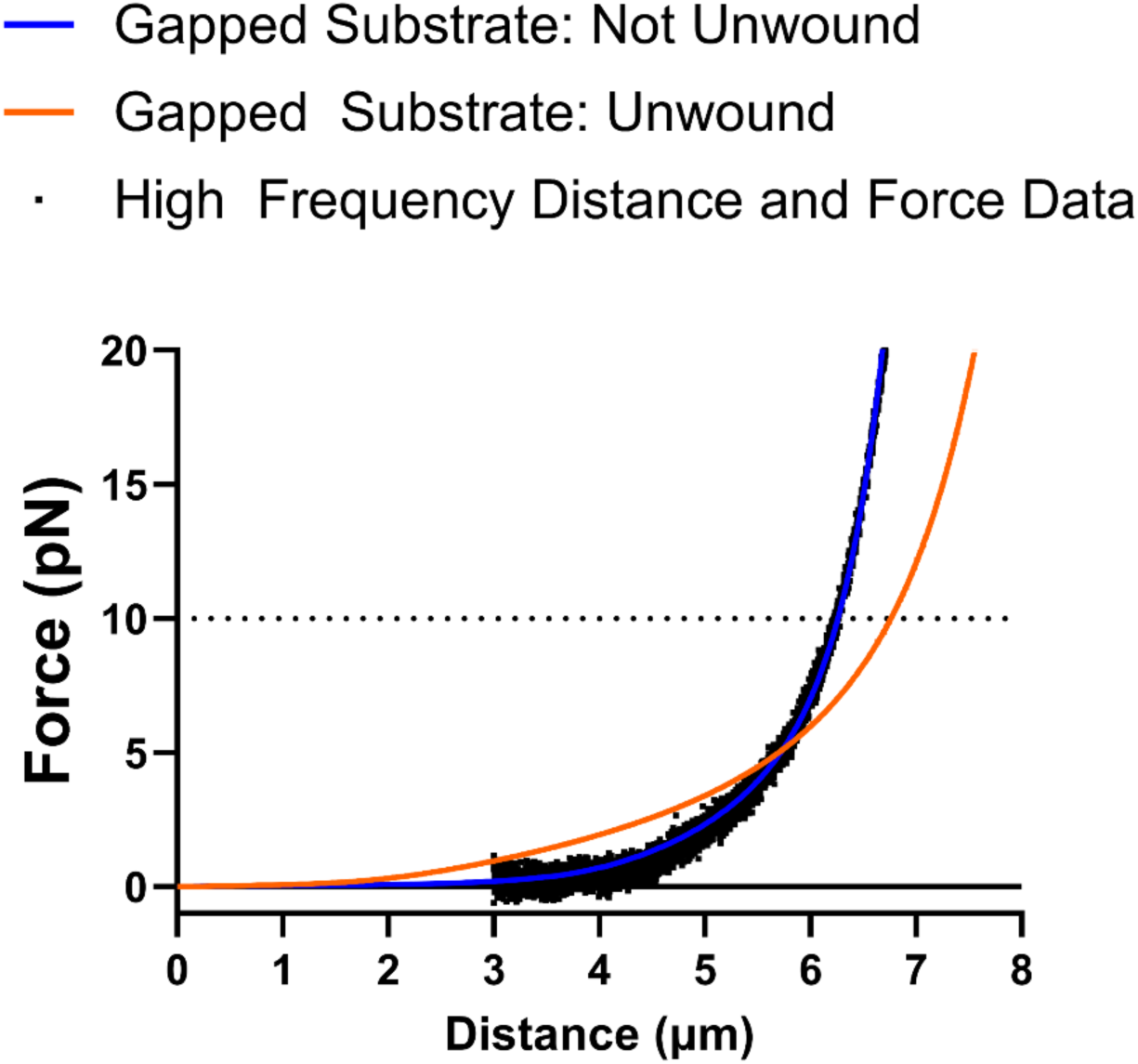
Determining DNA unwinding distance using the extensible Worm-Like Chain model. Simulated force-extension curves for the 5,005 nt gapped DNA used in the DNA unwinding experiments in Figures 5, 6 and 7. Blue curve is a simulation for the DNA consisting of 5,005 nts of ssDNA and 12,848 bp of dsDNA (both DNA handles) before DNA unwinding (simulated using **Equation 4**). Orange curve is the simulation for the fully unwound tether with 11,401 nts of ssDNA and one dsDNA handle of 6,452 bp. Black squares are data collected while extending the gapped DNA substrate and was down sampled to 200 Hz from 78,125 Hz by decimation. Data were collected in Single-Molecule Imaging Buffer in the absence of protein. The tether was extended twice before returning to the initial distance of 3 µm.

**Figure S10.**
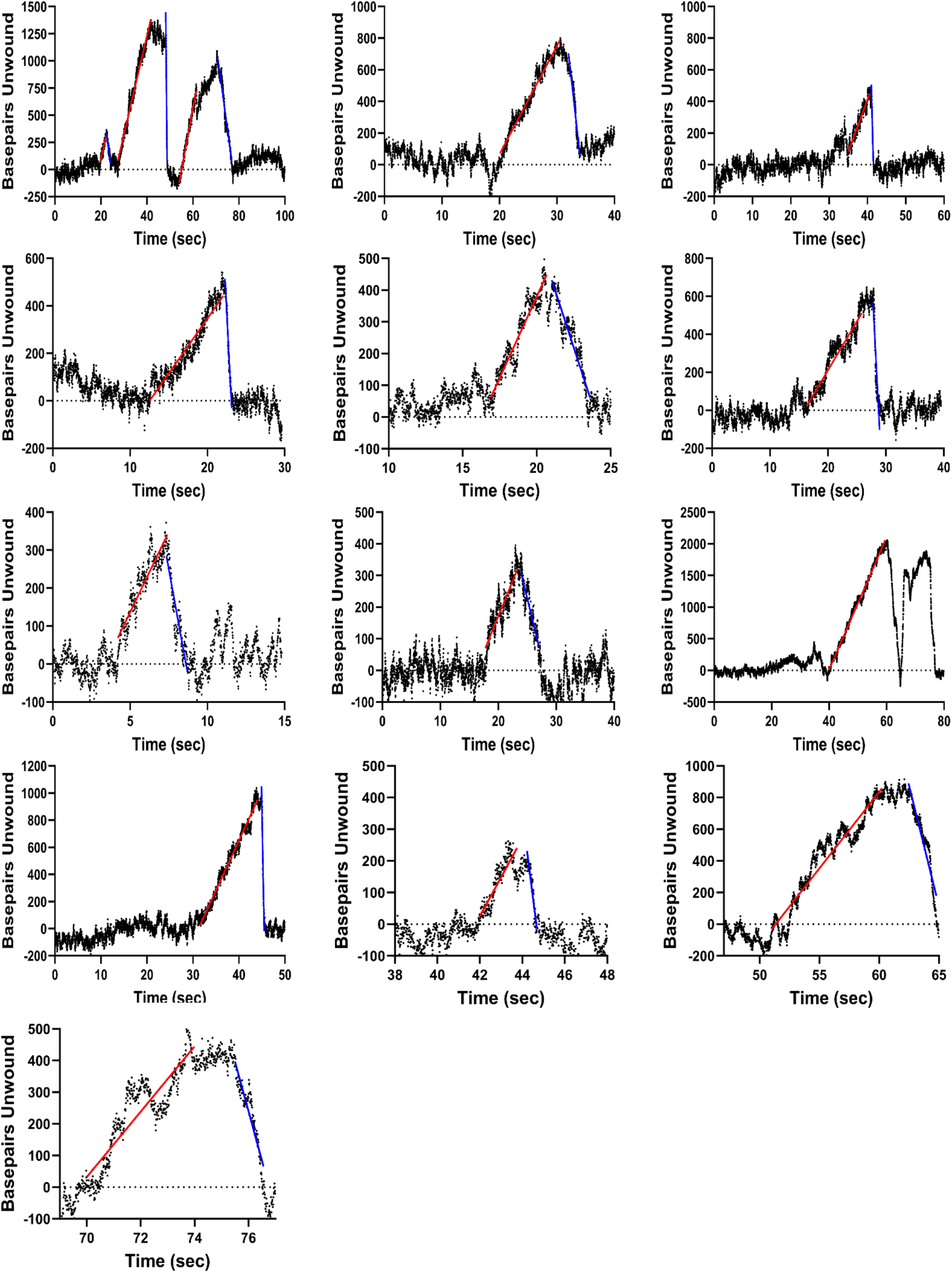
Individual DNA unwinding events for UvrD(R421C) dimers at 50 µM ATP associated with the data in Figure 5C & D. Each DNA unwinding event was obtained in Buffer T20-20, 50 μM ATP, 1 mM MgCl_2_ at 25°C. Red lines show the linear fits to DNA unwinding events and blue lines show linear fits to the rehybridization events.

**Figure S11.**
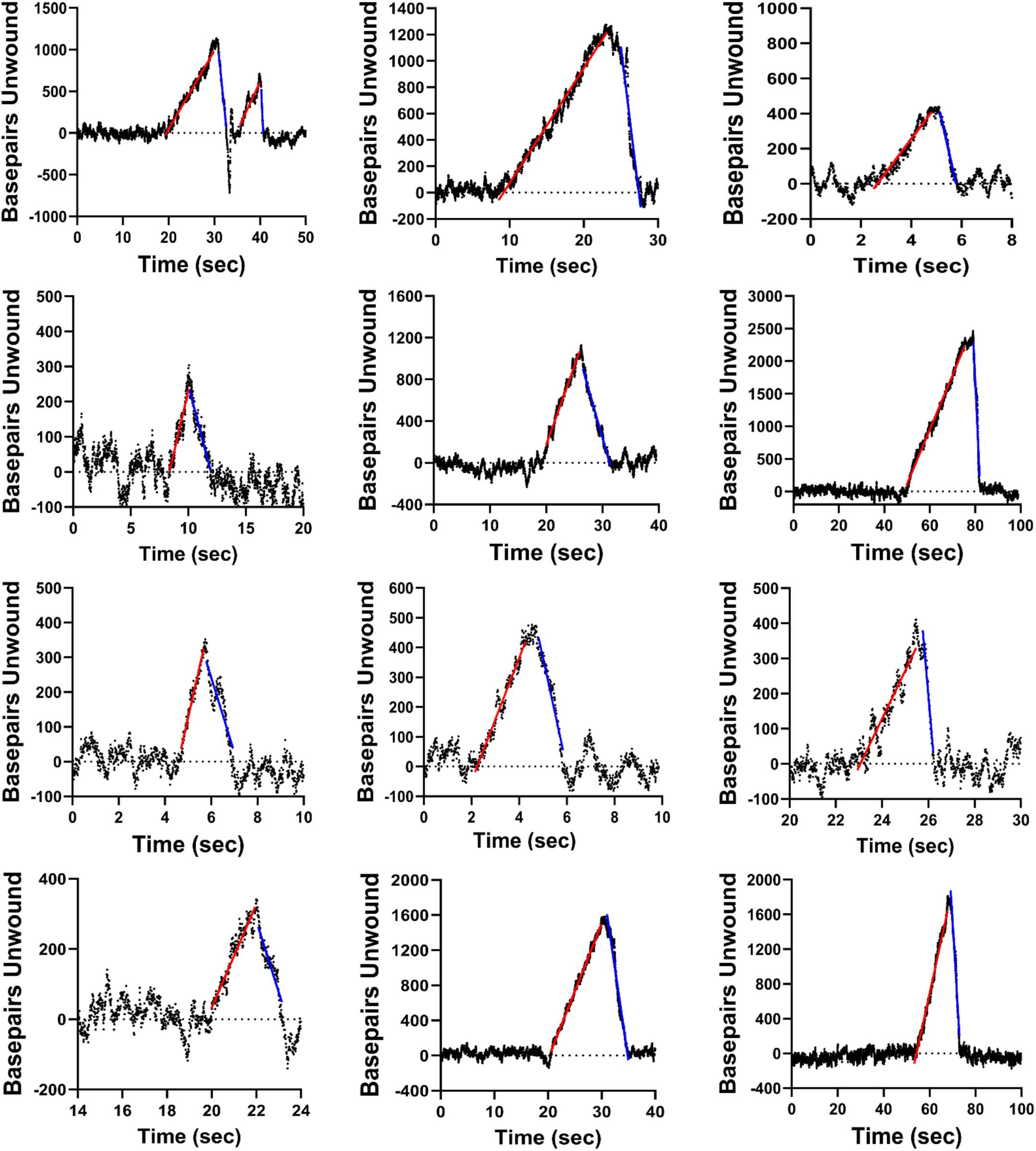
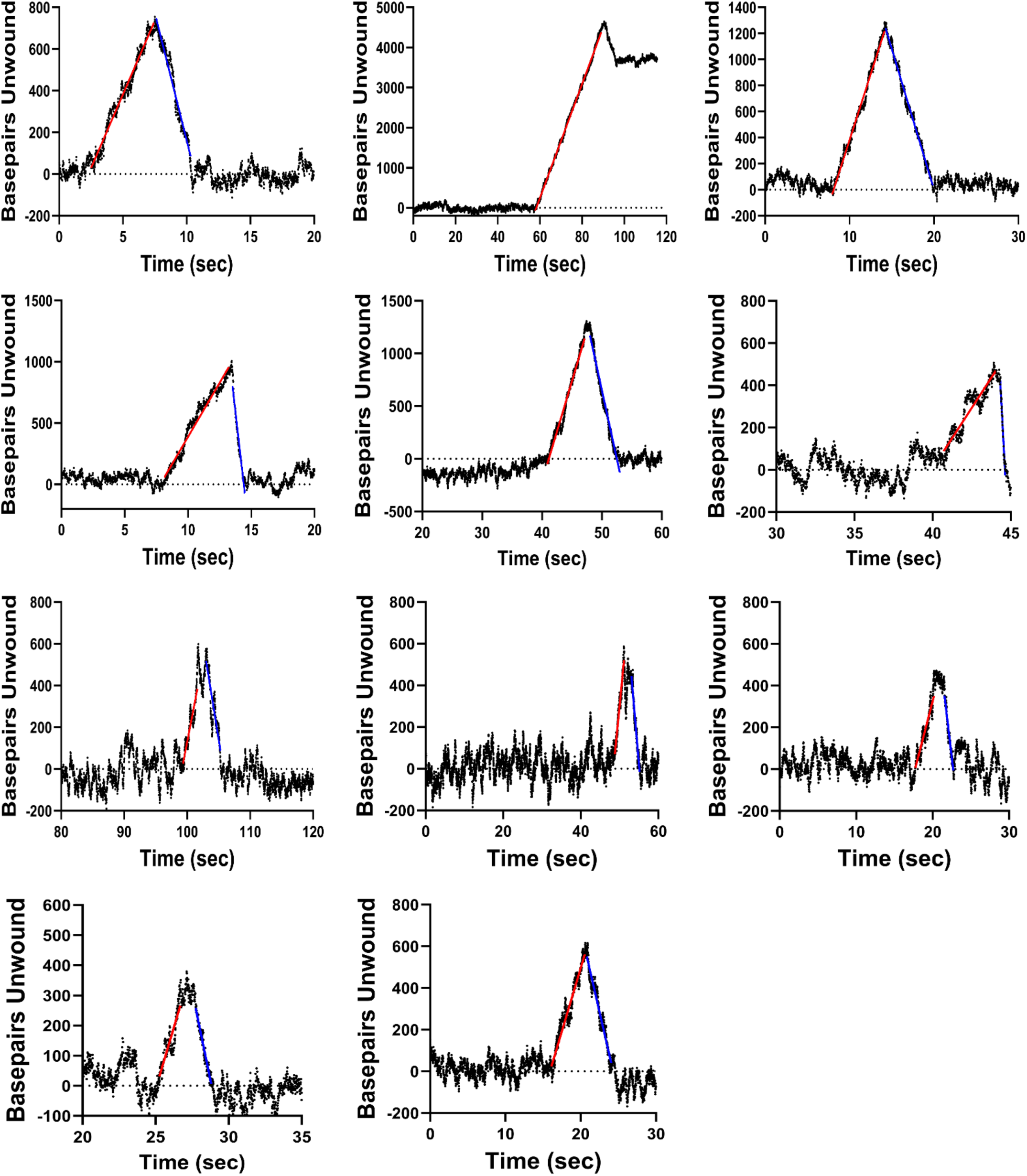
Individual DNA unwinding events for UvrD(R421C) dimers at 500 µM ATP in Figure 5C & E. Each DNA unwinding event was obtained in Buffer T20-20, 500 μM ATP, 1 mM MgCl_2_ at 25°C. Red lines show the linear fits to DNA unwinding events and blue lines show linear fits to the rehybridization events.

**Figure S12.**
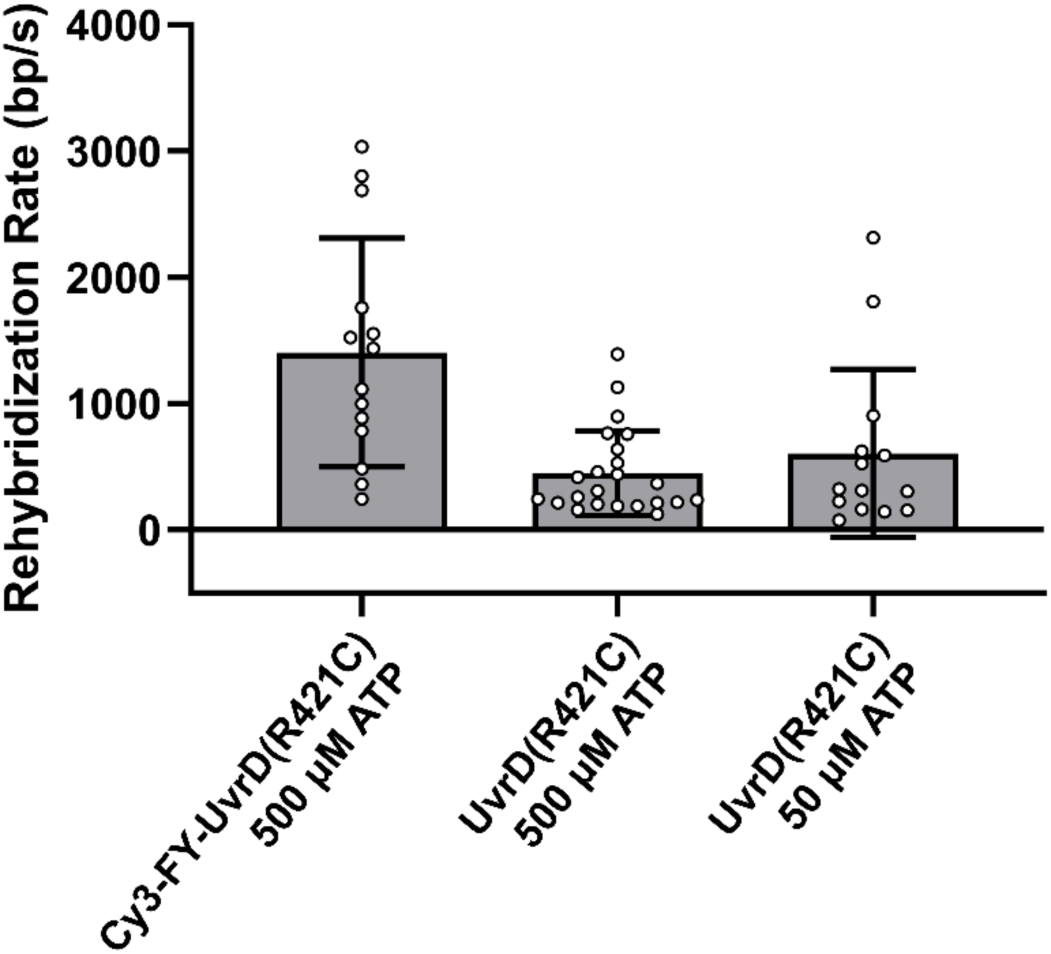
Rates of double-stranded DNA rehybridization. Rates of DNA rehybridization were determined by fitting the data in Figures S10, S11, & S13 (blue linear lines). DNA rehybridization rates are 1405±904 bp/s for Cy3-FY-UvrD(R421C) dimers collected in Single-Molecule Imaging Buffer (500 μM ATP) at 25°C, 450±336 bps/sec for UvrD(R421C) dimers in 500 μM ATP and 605±665 bps/sec for UvrD(R421C) dimers in 50 μM ATP in Buffer T20-20 and 1 mM MgCl_2_ at 25°C.

**Figure S13.**
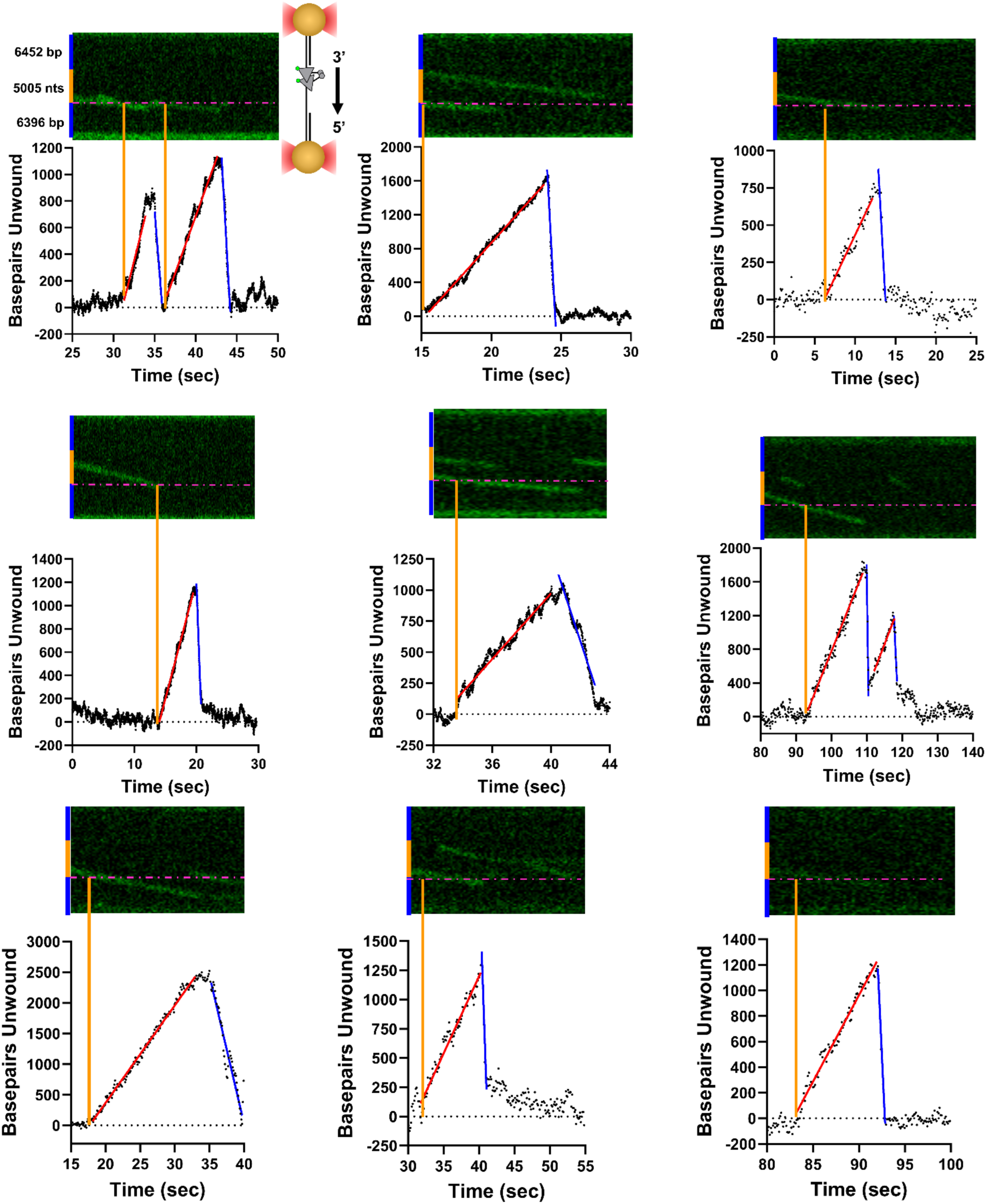
Individual DNA unwinding events for FY-UvrD(R421C) Cy3-labeled dimers (data in Figure 6D). Each DNA unwinding event was obtained in Single Molecule Imaging Buffer (500 μM ATP), 25 °C. Red lines show the linear fits to DNA unwinding events and blue lines show linear fits to the rehybridization events. Orange vertical lines indicate the point of initiation of DNA unwinding in both the kymograph (above) and DNA unwinding analysis obtained from the change in bead position (below).

## Notes

### Competing Interest Statement

The authors have declared no competing interest.

